# Evaluating the fitness of PA/I38T-substituted influenza A viruses with reduced baloxavir susceptibility in a competitive mixtures ferret model

**DOI:** 10.1101/2020.09.28.316620

**Authors:** Leo YY Lee, Jie Zhou, Paulina Koszalka, Rebecca Frise, Rubaiyea Farrukee, Keiko Baba, Shahjahan Miah, Takao Shishido, Monica Galiano, Takashi Hashimoto, Shinya Omoto, Takeki Uehara, Edin Mifsud, Neil Collinson, Klaus Kuhlbusch, Barry Clinch, Steffen Wildum, Wendy Barclay, Aeron C Hurt

**Author notes:** Department of Microbiology and Immunology, University of Melbourne, at the Peter Doherty Institute for Infection and Immunity, Parkville, Australia (Leo YY Lee).

## Abstract

Baloxavir is approved in several countries for the treatment of uncomplicated influenza in otherwise-healthy and high-risk patients. Treatment-emergent viruses with reduced susceptibility to baloxavir have been detected in clinical trials, but the likelihood of widespread occurrence depends on replication capacity and onward transmission. We evaluated the fitness of A/H3N2 and A/H1N1pdm09 viruses with the polymerase acidic I38T-variant conferring reduced susceptibility to baloxavir relative to wild-type (WT) viruses, using a competitive mixture ferret model, recombinant viruses and patient-derived virus isolates. The A/H3N2 I38T virus showed a reduction in within-host fitness but comparable between-host fitness to the WT virus, while the A/H1N1pdm09 I38T virus had broadly similar within-host fitness but substantially lower between-host fitness. Although I38T viruses replicate and transmit between ferrets, our data suggest that viruses with this amino acid substitution have lower fitness relative to WT and this relative fitness cost was greater in A/H1N1pdm09 viruses than in A/H3N2 viruses.

## Introduction

Antivirals are important for the treatment of influenza infections, particularly in high-risk individuals such as the immunocompromised and the elderly. Neuraminidase inhibitors (NAIs) such as oseltamivir are the current standard of care for treating influenza-related hospitalisations during seasonal epidemics (Heneghan et al., 2016), and are stockpiled in some countries as pandemic contingency (Patel et al., 2017). However, the high frequency of mutations during influenza virus replication combined with the selective pressure of antiviral treatment can lead to the emergence of viral variants with reduced antiviral susceptibility or resistance. The World Health Organization Global Influenza Surveillance and Response System regularly monitors circulating influenza viruses for antiviral resistance, as the prevalence of resistant variants circulating in the community is a key factor when considering which influenza antiviral drugs should be prescribed for clinical use. For example, M2 ion channel blockers are no longer prescribed to treat influenza A viruses, as nearly 100% of circulating influenza viruses contain an S31N amino acid substitution that confers resistance to these antiviral drugs (Hussain, Galvin, Haw, Nutsford, & Husain, 2017). Similarly, H275Y substitutions in viral neuraminidase (NA) can lead to reduced susceptibility of influenza viruses to oseltamivir and another NAI, peramivir (Li, Chan, & Lee, 2015). While localised clusters of H275Y oseltamivir-resistant A/H1N1pdm09 viruses have been detected (Hurt et al., 2012; Takashita et al., 2014), these variants are currently only detected at a frequency of <1% among circulating viruses (Takashita et al., 2020). There is an ongoing need for the development of antivirals that utilise novel mechanisms of action, such that multiple effective treatment options are available in the case that resistance towards an existing class of antiviral drugs becomes widespread.

Baloxavir (pro-drug: baloxavir marboxil; active form: baloxavir acid) has been approved for the treatment of uncomplicated influenza in otherwise-healthy and high-risk patients in several countries around the world. The influenza polymerase complex is a heterotrimer comprising the polymerase acidic (PA) and two polymerase basic (PB1 and PB2) proteins. Baloxavir targets the highly conserved cap-dependent endonuclease region of the PA to inhibit ‘cap-snatching’, a critical stage of influenza virus replication (Dias et al., 2009). Baloxavir has activity against influenza A, B, C and D viruses (Mishin et al., 2019; Noshi et al., 2018).

Human clinical trial data have shown that a single oral baloxavir dose alleviates influenza symptoms at least as effectively as a 5-day oral course of twice-daily oseltamivir, as well as suppressing viral shedding more rapidly in all patient populations examined (Baker et al., 2020; Hayden et al., 2018; Hirotsu et al., 2019; Ison et al., 2020). In 9.7% of otherwise-healthy patients treated with baloxavir in the phase 3 CAPSTONE-1 trial, viruses with amino acid substitutions at residue I38 of the PA protein were identified. The most frequent variant was I38T, that has subsequently been shown to exhibit reduced susceptibility to baloxavir in vitro (Hayden et al., 2018). I38T variants have emerged following baloxavir treatment in subsequent clinical trials, with the highest rate of 23.4% observed in paediatric patients (Dias et al., 2009; Hirotsu et al., 2019; Noshi et al., 2018). Although I38X variants are associated with up to 50-fold reduction in baloxavir susceptibility in vitro compared to wild type (WT) viruses (Omoto et al., 2018), clinical efficacy (time to alleviation of symptoms) is still observed in baloxavir-treated patients in whom these variants develop, presumably in part because resistance does not emerge until several days after treatment has begun (Uehara et al., 2019). Although based on small patient numbers, post-hoc analyses of the phase 3 CAPSTONE-1 trial suggested these variants can be associated with prolonged virus detection and uncommonly with symptom rebound (Uehara et al., 2019). However, analysis of the CAPSTONE-2 trial (patients at high risk of complications) showed the opposite effect, whereby I38X variants were associated with numerically faster resolution of symptoms than those without (Ison et al., 2020), suggesting further investigation into the clinical impact of I38X variants is required.

Variant viruses containing amino acid substitutions that reduce influenza antiviral susceptibility can also lower the efficiency of viral replication and transmission (Duan et al., 2010; Herlocher et al., 2002; Ives et al., 2002; Tu et al., 2017; Yen et al., 2005). The extent of these fitness impacts can vary depending on the specific substitution and the viral genetic background, and compensatory mutations can develop which restore viral fitness in resistant variants (Dong et al., 2015; Lackenby et al., 2018; Li et al., 2015; Simonsen et al., 2007). A notable example of this has been the oseltamivir- and peramivir-resistant A/H1N1 H275Y variant which, while initially showing impaired replicative capacity *in vitro* (Abed, Goyette, & Boivin, 2005; Ives et al., 2002), acquired several compensatory mutations resulting in the capacity for a fit H275Y variant to arise and spread globally in 2008 (Dharan et al., 2009; Hurt et al., 2009; Meijer et al., 2009). The resistant virus was outcompeted from circulation by the newly emergent oseltamivir-sensitive pandemic A/H1N1pdm09 strain in 2009.

Although treatment-emergent viruses with reduced susceptibility to baloxavir have been detected in clinical trials, the public health risk and likelihood of their widespread occurrence depends on their capacity to replicate and transmit compared with WT viruses. To evaluate the intrinsic fitness of I38T-variant A/H1N1pdm09 or A/H3N2 viruses with those that have reduced baloxavir susceptibility, we used in vitro assays and in vivo ferret models to compare their replication and transmission efficiency relative to baloxavir-sensitive (WT) viruses, using both recombinant viruses and patient-derived clinical virus isolates from human trials.

## Results

### Baloxavir susceptibility of I38T-variants from post-treatment clinical specimens

Pre- and post-treatment clinical isolate viruses were tested by plaque reduction assay to assess the baloxavir susceptibility of treatment-emergent variant viruses harbouring the I38T substitution. The mean baloxavir EC_50_ of viruses from pre-treatment samples (WT) were 1.1–1.5 and 0.31–0.69 ng/mL in A/H1N1pdm09 and A/H3N2 viruses, respectively, whereas the corresponding baloxavir EC_50_ of the post-treatment viruses (I38T) were 82–87 and 36– 53 ng/mL, respectively. Therefore, A/H1N1pdm09 and A/H3N2 I38T-variants displayed 58– 75- and 77–155-fold reductions in baloxavir susceptibility compared with their respective WT viruses, similar to levels reported previously (Imai et al., 2019; Noshi et al., 2018; Uehara et al., 2019).

### In vitro comparison of WT and I38T-variant replication kinetics

The in vitro replicative capacity of pre-treatment WT and matched post-treatment I38T-substituted clinical isolates was compared in MucilAir™ human nasal epithelial cells (*Figure 1A*). For the two A/H1N1pdm09 virus pairs, we observed transient reductions of viral titres in the I38T viruses compared to WT at 48 hours in sample 2PQ003 (p<0.05) and at 72 hours (p<0.05) in sample 2HB001. No significant differences were observed in virus titres for A/H3N2 clinical isolates, indicating that in vitro growth kinetics of the A/H3N2 I38T-variants were comparable to the WT viruses.

**Figure 1.**
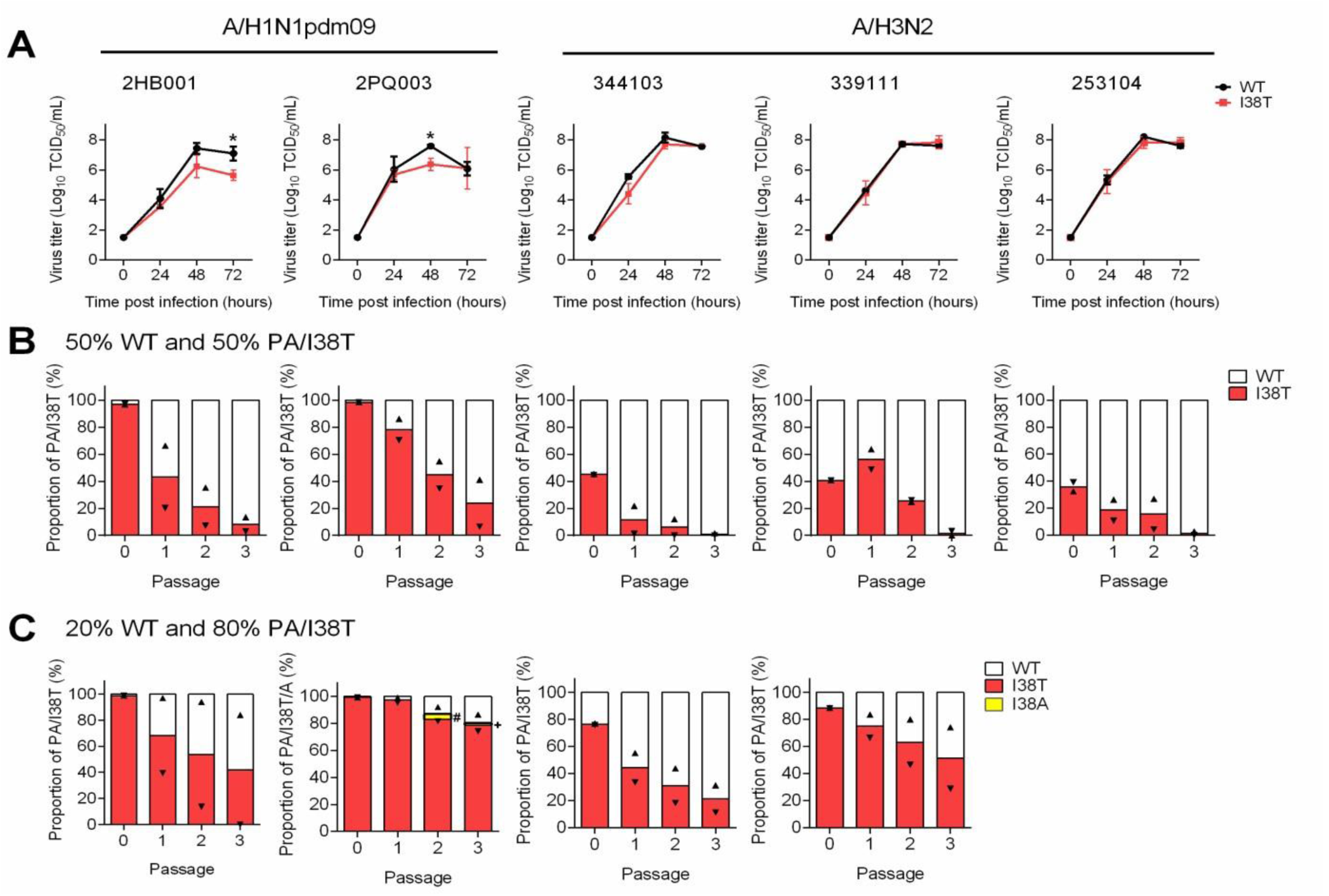
Replicative capacity and competitive fitness of influenza PA/I38T substituted A/H1N1pdm09 and A/H3N2 viruses isolated from clinical samples. (A) MucilAir™ human nasal epithelial cells were infected with influenza clinical isolates at an MOI of 0.001 TCID_50_/cell. Culture fluids were collected at 0, 24, 48 and 72 hours postinfection and viral titres were determined in MDCK-SIAT1 cells. The LLOQ for virus titre was 1.57 log_10_ TCID_50_/mL and the viral titre lower than LLOQ was defined as 1.50 log_10_ TCID_50_/mL. Statistical comparisons of virus titres between WT and viruses with I38T at each time-point were conducted using Welch’s *t*-test at the 0.05 significance level (*p<0.05). Data represent mean ± SD of triplicate experiments. (B and C) MucilAir™ cells were co-infected with WT and I38T-substituted clinical isolates at (B) 50:50 or (C) 20:80 ratio based on viral titres (TCID_50_) at an MOI of 0.002. At 48 hours post-infection, the culture fluids were collected and serially passaged three times at an MOI of 0.001. Each culture fluid was subjected to NGS to determine the population of amino acids at position 38 in the PA subunit. The lower detection limit for calling variant viruses was 1% and variant population less than the lower detection limit was set as 0 for the calculation. The bar and the triangles indicate the mean and the individual values of duplicate experiments, respectively. The symbols (# and +) indicate 3.8% and 1.5% of I38A proportion. LLOQ, lower limit of quantitation; MOI, multiplicity of infection; NGS, next generation sequencing; PA, polymerase acid; SD, standard deviation; TCID_50_, 50% tissue culture infectious dose; WT, wild type.

A competitive mixtures experiment design was used to further investigate the effect of I38T on in vitro replication fitness (*Figure 1B and C*). Following inoculation of virus mixtures, we evaluated variant viral fitness based on the change in the percentage of I38T viruses (%I38T) in the viral population over three successive passages harvested from the infected cell supernatants. For all virus pairs tested, the proportion of the I38T virus replicating in MucilAir™ cells decreased with each passage, regardless of virus subtype or initial ratio. Interestingly, a new I38A variant virus was detected by next generation sequencing (NGS) in a #2PQ003 co-infection group at P2 and P3 (*Figure 1C*), however this subpopulation was transient and remained at low abundance. Compared to the A/H3N2 clinical isolates, the ratio of WT%:I38T% determined by NGS for A/H1N1pdm09 clinical isolate pairs did not correlate well with the intended mixture proportions based on infectious virus titre (used for preparing the P0 material), which may be due to the presence of non-replicating virions present in the viral samples (*Figure 1B and C*). Despite these mixtures consisting of ∼99% I38T viral copies based on NGS, mixed virus populations similar to the expected proportions were established at 48 hours following inoculation (P1, *Figure 1B and C*), and we identified that the A/H1N1pdm09 I38T-variant population declined compared to WT with each successive passage. In addition to these human cell experiments, we performed a competitive mixtures experiment in MDCK cells using recombinant WT and I38X-variant virus pairs. As with previous experiments, WT viruses to exhibit a replication advantage over I38X-variants (*Figure 1 – figure supplement 1*).

The A/H3N2 clinical isolate pair #344103 (WT and I38T) and A/H1N1pdm09 clinical isolate pair #2HB001 (WT and I38T) were selected to investigate further for viral fitness in the ferret infection model (*Figure 1 – figure supplement 2*).

### Replication and direct contact transmission of WT/I38T clinical isolate virus pairs in a competitive fitness ferret model (Melbourne)

#### A/H3N2 I38T fitness

In the Melbourne laboratory, the in vivo replication and transmission capacity of the post-treatment A/H3N2 I38T isolate was compared to the corresponding pre-treatment WT isolate in ferret transmission chains. Infectious virus was detected in nasal washes of each recipient ferret (RF) in transmission chains initiated with pure populations of either the WT or I38T virus, indicating that the treatment-emergent I38T-variant was capable of at least three successive transmissions in a direct contact model (*Figure 2A*). Pyrosequencing analysis showed that the I38T-variant population was maintained over three generations of transmission without substantial reversion to WT (*Figure 2 – figure supplement 1*). To compare replication kinetics of A/H3N2 I38T virus with WT virus in vivo, the area under the curve (AUC) of infectious viral titres in the nasal washes of RF1-RF3 (n = 3) was determined over the course of infection (*Figure 2A*). Similar to the in vitro experiments (*Figure 1A*), overall RF virus replication kinetics (AUC of the 50% tissue culture infectious dose [TCID_50_] ± standard deviation [SD]) for the A/H3N2 WT virus (18.42 ± 7.29) were comparable to the I38T virus (17.58 ± 2.75, non-significant).

**Figure 2.**
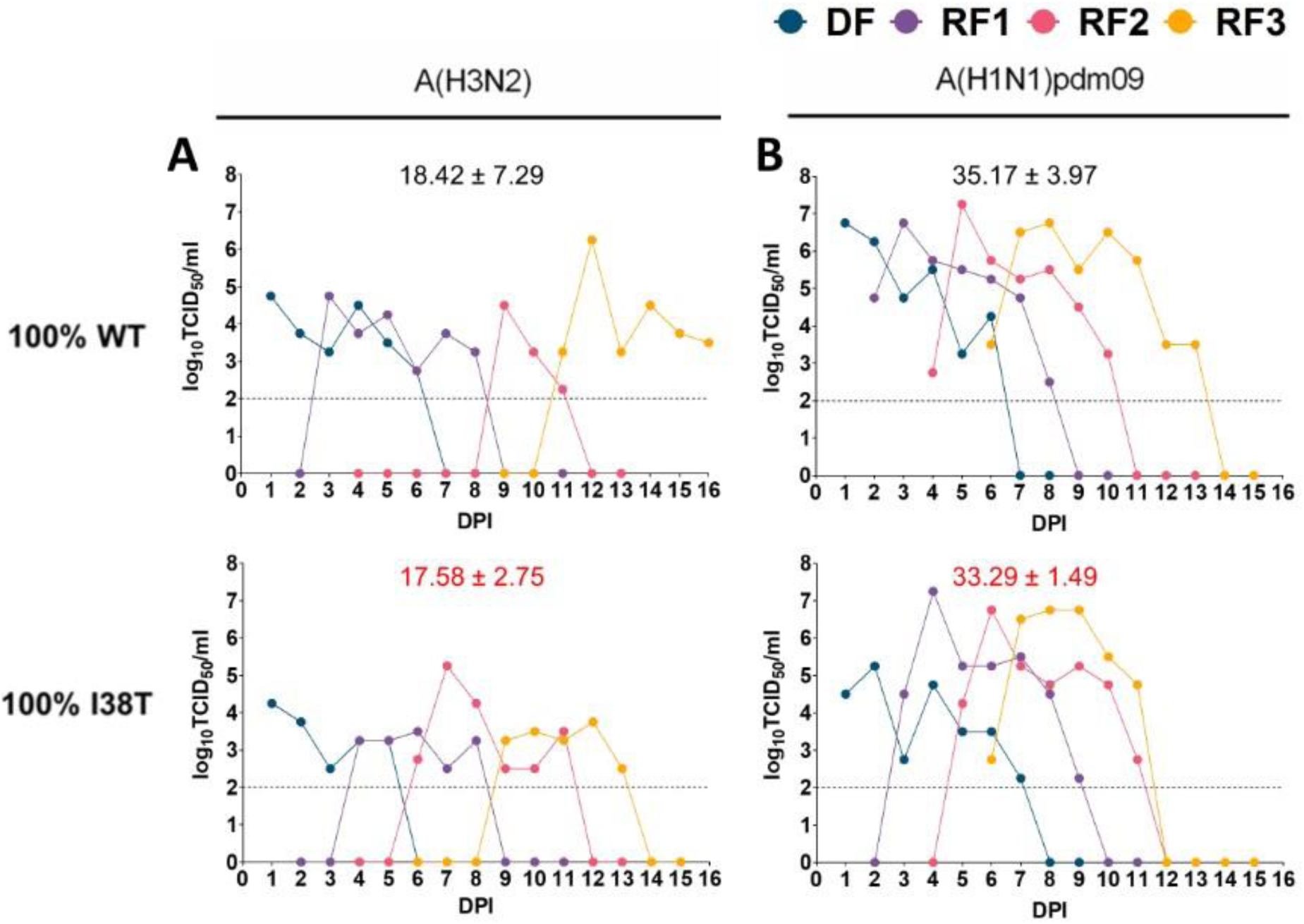
Infectious viral titres (TCID_50_) in nasal washes of direct contact recipient ferrets 1-3 (RF1-RF3) from pure (100%) WT and pure PA/I38T infection groups. (A) A/H3N2 clinical isolates; (B) A/H1N1pdm09 clinical isolates. Virus replication kinetics (TCID_50_) in ferrets infected by direct contact transmission (DF and RF1-RF3) using pure populations of WT or I38T-variant virus for (A) A/H3N2 or (B) A/H1N1pdm09 subtypes. Points represent infectious titres (log_10_ TCID_50_/mL) in individual ferret nasal washes from each DPI of DF. The mean AUC of infectious virus shedding (AUC TCID_50_ ± SD) for RF1-RF3 in each group is labelled (black text, WT; red text, I38T). AUC, area under the curve; DF, donor ferret; DPI, days post inoculation; RF, recipient ferret; SD, standard deviation; TCID_50_, 50% tissue culture infectious dose; WT, wild type.

In parallel, transmission chains were initiated with competitive virus mixtures in donor ferrets (DFs) inoculated with different ratios of A/H3N2 WT and I38T viruses based on infectious titres. Our aim was to establish infections in DFs with WT and I38T-variant viruses present at a range of different ratios, allowing us to evaluate changes over time for the proportion of I38T-variant population in infected ferrets and the differences in variant population following a transmission event to a recipient ferret. Pyrosequencing of viral RNA shedding in nasal wash samples showed that the mixed infections established in DFs at 1 day post inoculation (DPI) were similar to the intended infectious virus ratios in the inoculum mixture (*Figure 3*). To evaluate the relative within-host fitness of the I38T-variant, in each individual ferret the %I38T population on the final day of nasal washing was compared to the population present on the first day of detection by pyrosequencing. We observed that the proportion of A/H3N2 I38T either remained stable (in DFs, with some decreases observed in [WT:I38T] 50:50 and 20:80 chains) or decreased over time (in RF1s) (*Figure 3*). The study of within host fitness of A/H3N2 and I38T viruses was limited when the viral populations established in RF2-RF3 were nearly pure (*Figure 3*), and therefore this resulted in limited competition of WT and I38T virus. The relative changes in A/H3N2 I38T proportion that occurred in each animal is summarised in *Figure 4A*; overall there was a modest reduction in the within-host fitness of the A/H3N2 I38T-variant. Transmission or between-host fitness was evaluated according to the change in variant virus population in the first pyrosequencing detection (post infection) in a RF nasal wash, compared to the population detected in the cognate DF on the previous day. Between-host fitness was similar in the I38T-variant and the WT A/H3N2 virus at each direct contact transmission event (*Figure 4B*). Some shifts (both increase and decrease) in %I38T were observed in the first generation of transmission (purple points), but apparent transmission bottlenecks of near pure WT (left extreme of x-axis) or I38T (right extreme of x-axis) had developed in the second (pink points) and third (yellow points) generations such that competition between I38T and WT virus populations was effectively absent in RF2-RF3, and minimal changes in proportion were observed (*Figure 4B*).

**Figure 3.**
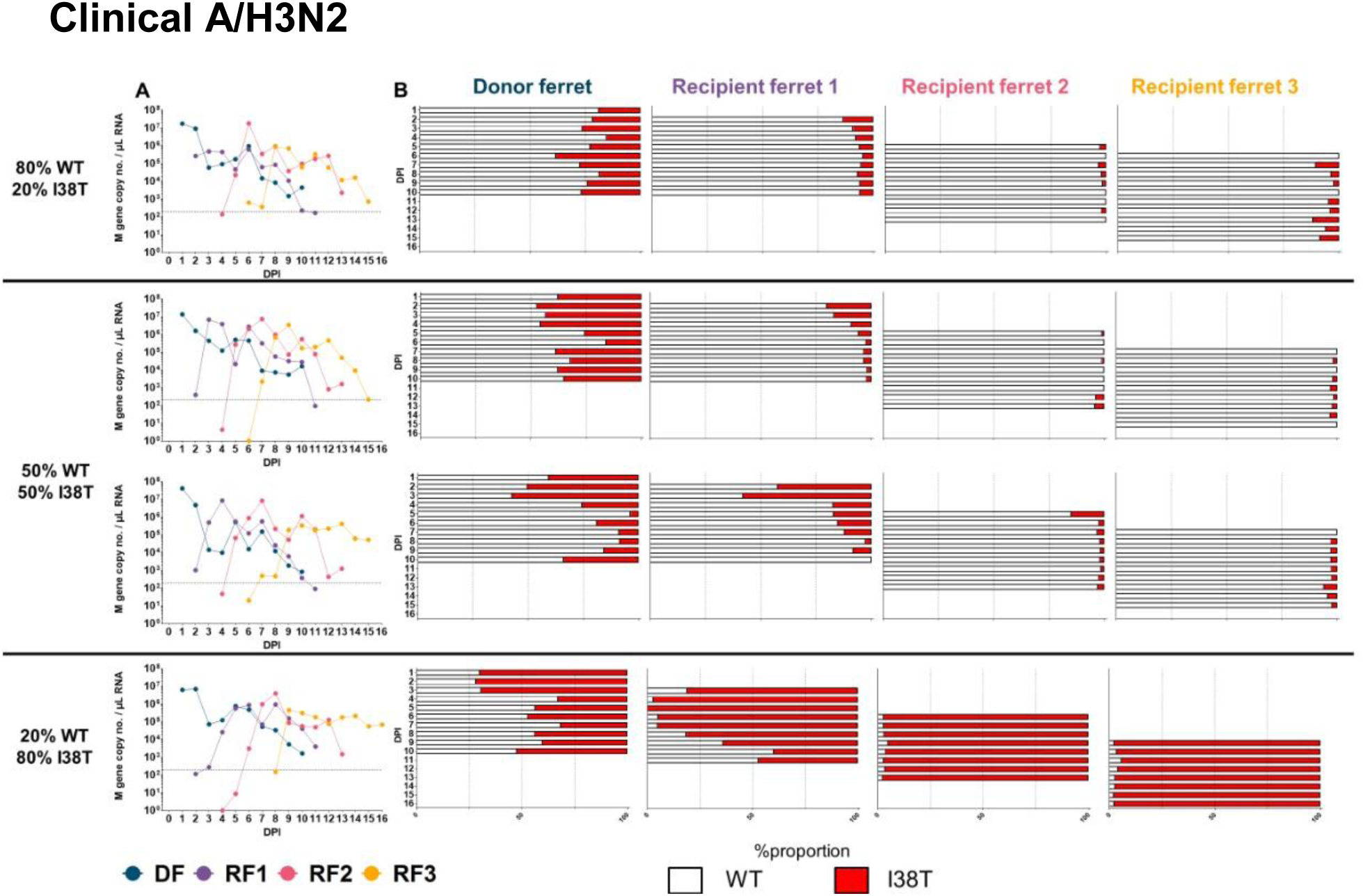
A) qPCR curves for A/H3N2 viral RNA and B) WT%:I38T% of viruses in ferret nasal washes. A/H3N2 DFs were infected (intranasal inoculation) with virus mixtures of WT and I38T at a total infectious dose of 10^5^ TCID_50_/ 500 μL. Daily nasal washes were collected from all animals for 10 consecutive days following inoculation/exposure, or until endpoint. RF1 was exposed to the DF by cohousing at 1 DPI. Naïve recipients were cohoused sequentially with the previous recipient (1:1) 1 day following detection of influenza A in RF nasal wash (Ct <35). (A) Viral RNA was quantified by number of M gene copies (influenza A)/μL of total RNA. (B) The relative proportion of virus encoding WT PA (white bars) and I38T (red bars) in each nasal wash sample was determined by pyrosequencing. DF, donor ferret; DPI, days post inoculation; RF, recipient ferret; TCID_50_, 50% tissue culture infectious dose; WT, wild type.

**Figure 4.**
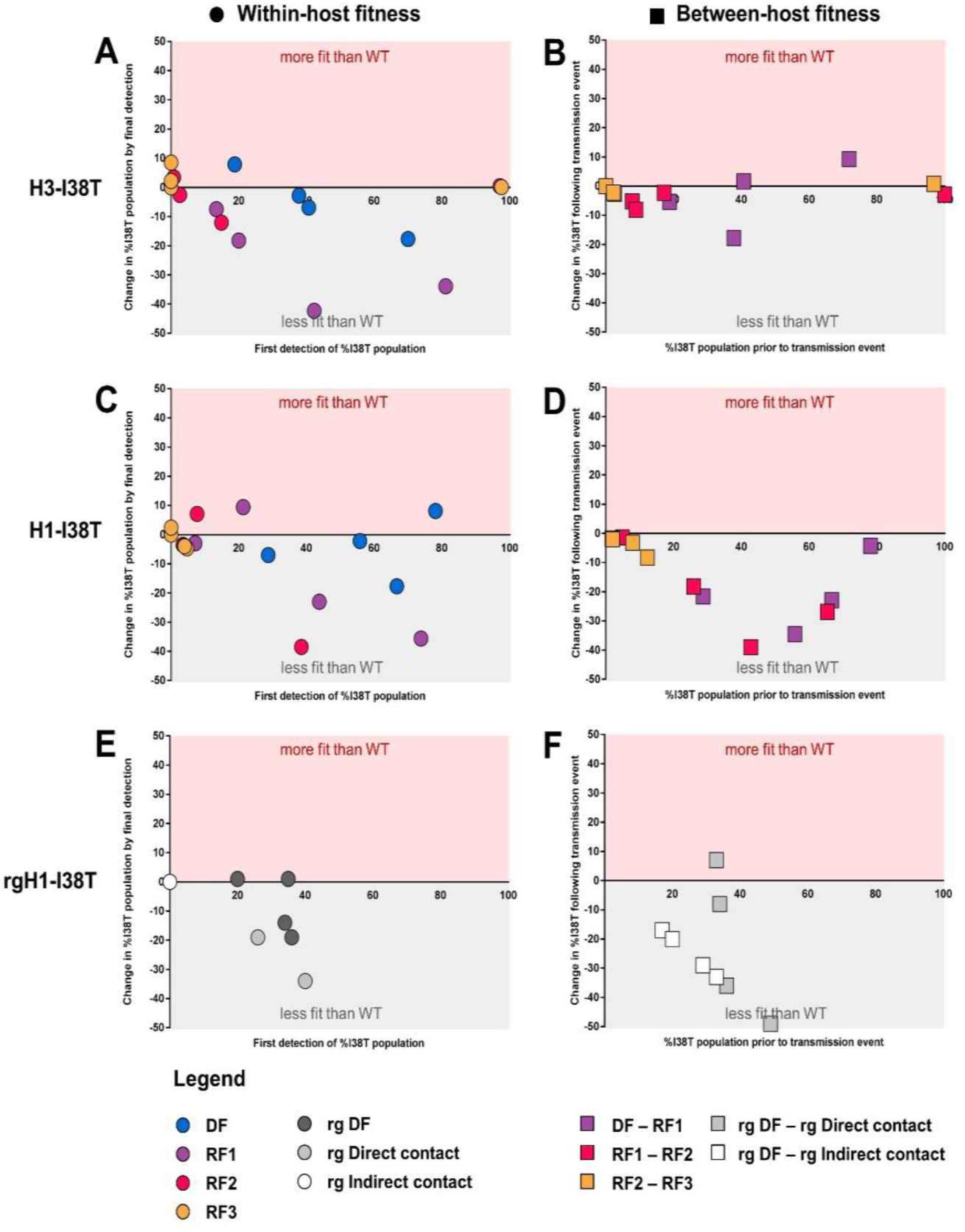
Visualisations of estimated within-host fitness and between-host fitness for PA/I38T-variant viruses. Within-host, and between-host transmission fitness visualisation for I38T-substituted clinical isolates (A–D) and recombinant viruses (E and F). (A, C and E) Within-host fitness graph Y-axis plots the change in %I38T population from first valid pyrosequencing/NGS detection (on X-axis) compared to last detection in each individual ferret (coloured circles represent DF: blue; RF1: purple; RF2: pink; RF3; yellow; rg DF: charcoal; rg direct contact: grey; rg indirect contact: white). Points in red shaded area indicate %I38T increased over time, grey shaded area indicates %I38T reduced over time. (B, D and F) Between-host fitness graph Y-axis plots the change in %I38T population for each transmission event measured using the first valid pyrosequencing/NGS measurement of %I38T in the recipient compared to the %I38T in the respective donor on the previous day (donor value is the X-axis). Coloured squares represent generations of transmission (1st, DF-RF1: purple; 2nd, RF1-RF2: pink; 3rd, RF2-RF3: yellow) or route of transmission (grey: direct contact; white: indirect contact). Points in red shaded area indicate %I38T increased following transmission, grey shaded area indicates %I38T reduced following transmission. DF, donor ferret, NGS, next generation sequencing; RF, recipient ferret.

Overall, the populations of A/H3N2 I38T-variant diminished such that they were essentially lost by RF2 in the three transmission chains that were initiated with either 20% or 50% of the variant. However, in the transmission chain initiated with 80% A/H3N2 I38T, the WT virus was outcompeted, likely to be due to the impact of a small transmission bottleneck in RF1 when the variant virus was still dominant in the viral mixture at 5 DPI (*Figure 3*). Therefore, across the two measured components of viral fitness, we identified that the A/H3N2 I38T virus had a modest reduction in within-host fitness, but comparable between-host fitness to that of the WT virus.

#### A/H1N1pdm09 I38T fitness

As observed in the A/H3N2 clinical isolate pair, infection with a pure population of the A/H1N1pmd09 I38T virus was capable of three successive direct contact transmissions in the ferret model (*Figure 2B*), and that the I38T substitution was maintained without reversion to WT (*Figure 2 – figure supplement 2*). Similar to the in vitro data for 2HB001 (*Figure 1A*), we measured a transient reduction in mean viral titre of A/H1N1pdm09 I38T recipients compared to WT at 24 hours post-exposure (p≤0.05). However, the overall infectious virus kinetics in ferrets that were infected with the A/H1N1pdm09 I38T virus (n = 3) (AUC 33.29 ± 1.49) was not significantly different to those infected with the WT virus (35.17 ± 3.97, non-significant) (*Figure 2B*).

In transmission chains initiated with mixtures of A/H1N1pdm09 WT and I38T-variant viruses, competitive co-infections were established at 1 DPI in DFs at similar proportions to the infectious virus ratio in the inoculum (*Figure 5*). In terms of within-host fitness, the proportion of A/H1N1pdm09 I38T remained relatively stable over time, except in the WT:I38T 20:80 chain, where some reductions were observed (*Figure 5*). This is further reflected in the summary figure where the majority of within-host measurements show little or no change in viral proportion, but in a small number of ferrets we observed a loss of the I38T-variant over time (*Figure 4C*). The I38T-variant population decreased following every direct contact transmission event, indicating the between-host fitness of A/H1N1pdm09 I38T was clearly compromised compared to WT (*Figure 4D*). The largest reductions in %I38T were observed in the first (purple points) and second (pink points) generations (*Figure 4D*), and by the endpoint, the I38T-variant proportion had become a minority proportion in all transmission chains (*Figure 5B*). Therefore, across the two measured components of viral fitness, we identified that the A/H1N1 I38T virus had broadly similar within-host fitness, but substantially lower transmission fitness compared to the WT virus.

**Figure 5.**
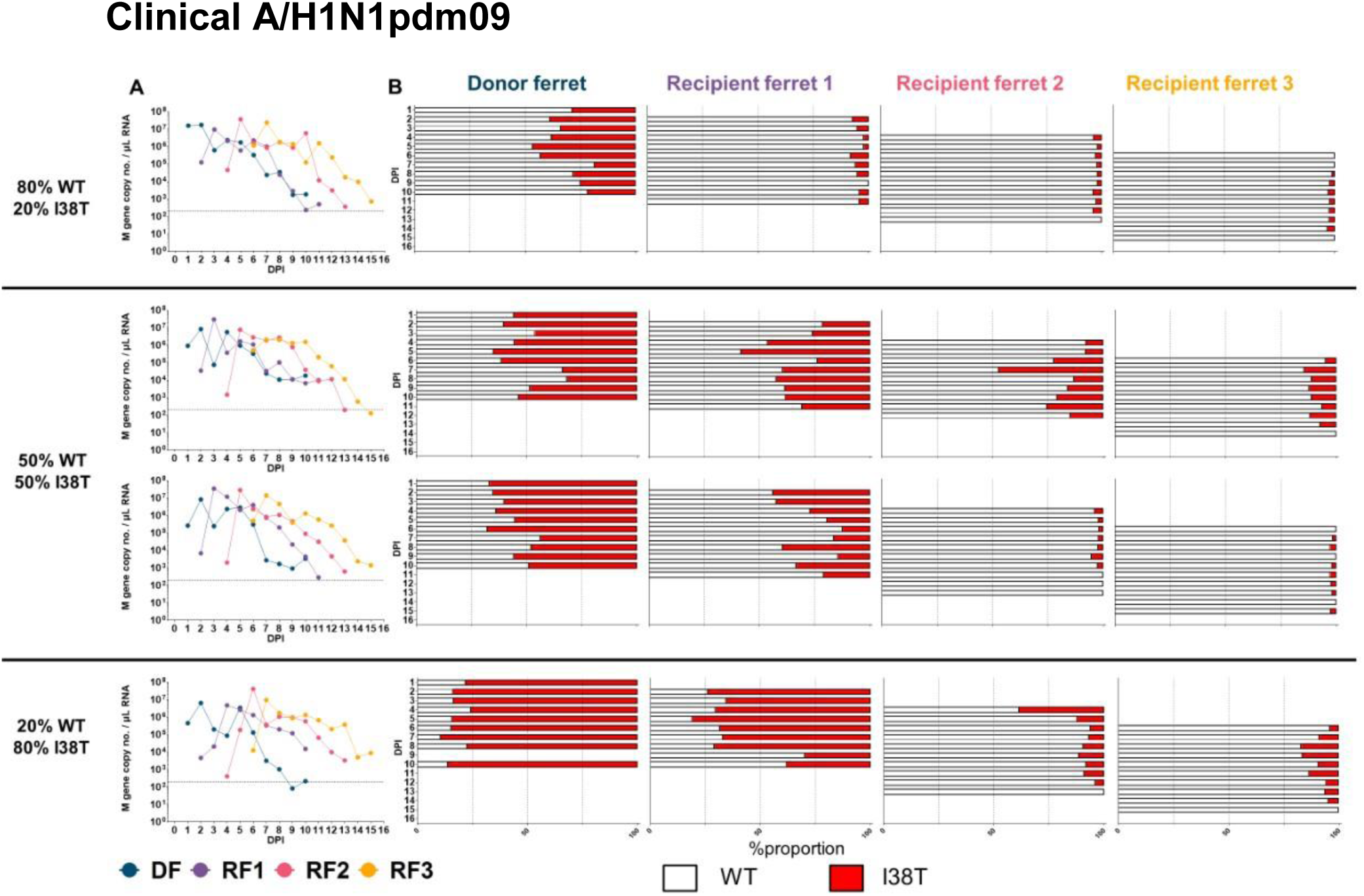
qPCR curves for A/H1N1pdm09 viral RNA and B) WT%:I38T% of viruses in ferret nasal washes. A/H1N1pdm09 DFs were infected (intranasal inoculation) with virus mixtures of WT and I38T at a total infectious dose of 10^4^ TCID_50_ / 500 μL. Daily nasal washes were collected from all animals for 10 consecutive days following inoculation/exposure, or until endpoint. RF1 was exposed to the DF by cohousing at 1 DPI. Naïve recipients were cohoused sequentially with the previous recipient (1:1) 1 day following detection of influenza A in RF nasal wash (Ct <35). (A) Viral RNA was quantified by number of M gene copies (influenza A) per μL of total RNA. (B) The relative proportion of virus encoding WT PA (white bars) and I38T (red bars) in each nasal wash sample was determined by pyrosequencing. DF, donor ferret; DPI, days post inoculation; PA, polymerase acid; RF, recipient ferret; TCID_50_, 50% tissue culture infectious dose; WT, wild type.

### Replication and direct/indirect contact transmission of WT/I38T reverse genetics A/H1N1pdm09 viruses in competitive fitness ferret model (London)

In the London laboratory, we tested the fitness impact of the I38T amino acid substitution in a reverse genetics-derived A/H1N1pdm09 virus using both direct and indirect transmission routes. DFs were inoculated with a 50:50 mixture before exposure (1:1:1) to both a direct contact recipient and an indirect contact recipient. NGS was used to determine the ratio of WT:I38T virus populations in ferret nasal washes, and showed that mixed infections were established in DFs at 1 DPI (*Figure 6*).

**Figure 6.**
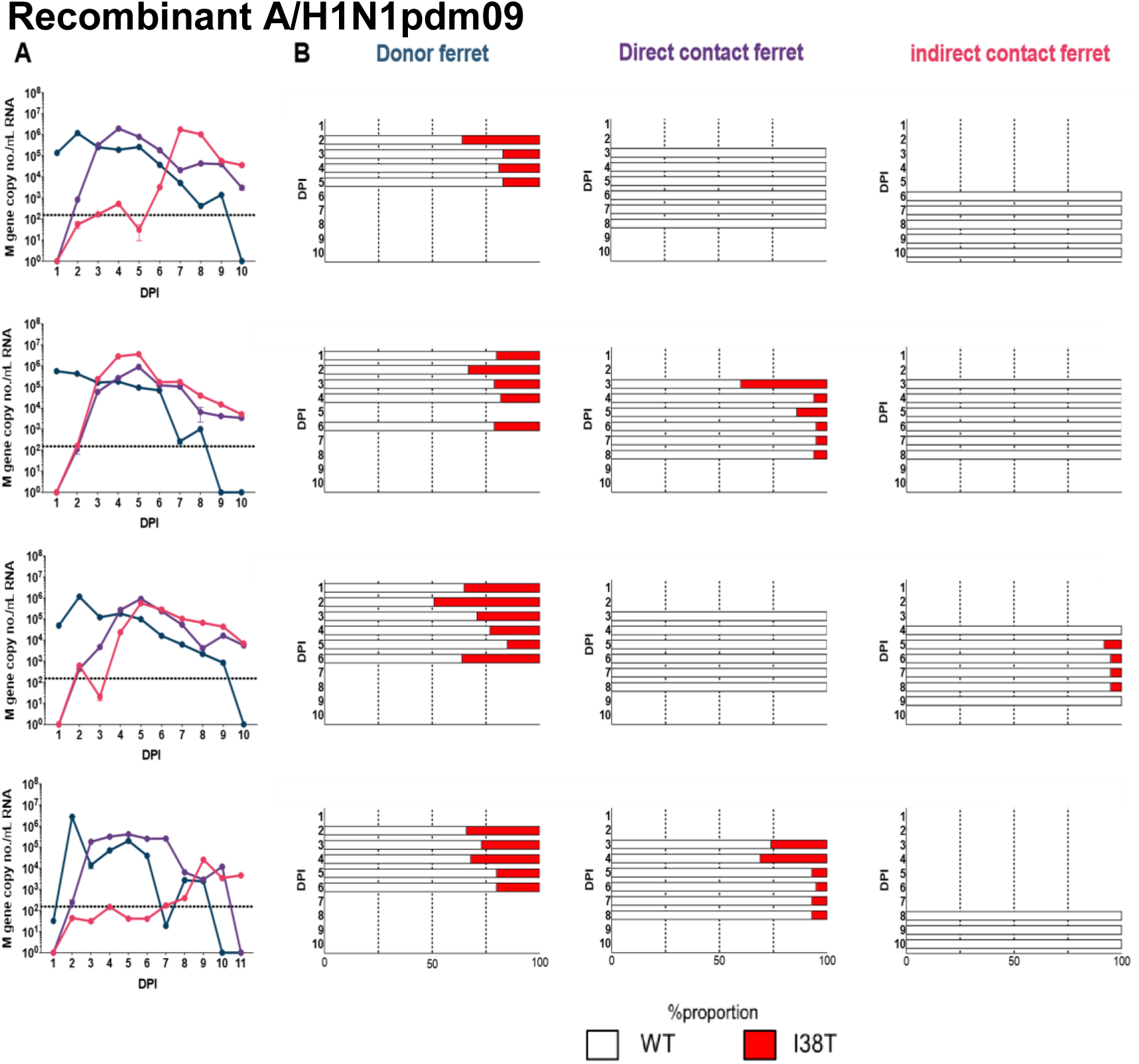
Change in WT/I38T virus proportion in nasal washes of donors compared to DC or IC recipients. Donor ferrets were infected (intranasal inoculation) with a recombinant A/H1N1pdm09 virus pair using 50:50 WT%: I38T% mixture based on infectious virus titre, at a total infectious dose of 10^4^ PFU. Naïve recipient ferrets were exposed to infected donors at 1 DPI by cohousing in the same cage (direct contact), or by housing in an adjacent cage separated by a respiratory droplet-permeable interface (indirect contact) (1:1:1 ratio). The relative proportion of virus encoding WT PA (white bars) and I38T (red bars) in each nasal wash sample was determined by next generation sequencing. DPI, days post inoculation; inoc, inoculation; PA, polymerase acid; WT, wild type.

In terms of within-host fitness, we observed that the I38T-variant population was maintained at a similar proportion in DF throughout the course of infection, but diminished rapidly in the direct contact ferrets where a mixed infection had been established by transmission (*Figure 6* and *4E*). In terms of between-host fitness the I38T-variant virus did not transmit by direct contact in two of four occasions (*Figure 6*), but the proportion remained relatively stable where transmission did occur (*Figure 4F*). Among indirect contact recipients, the I38T-variant virus failed to transmit in three of four occasions, and in the one of four recipients where it was detected it was at a very low abundance (<10%, near limit of detection) (*Figure 6*). These data support the findings from the A/H1N1pdm09 clinical isolates derived from treated patients, indicating that between-host fitness is substantially compromised by the I38T substitution. Furthermore, the results suggest that the transmission fitness of the I38T virus is lower in the indirect transmission model compared to the direct contact model (*Figure 4F*).

## Discussion

The detection of variant influenza viruses carrying the PA/I38T amino acid substitution in baloxavir-treated patients raises the concern that such viruses with reduced drug susceptibility could achieve sustainable transmission and circulate in the community, thereby potentially compromising the effectiveness of baloxavir usage. The present study provides evidence that the I38T substitution has some impact on aspects of the intrinsic viral fitness of both A/H3N2 and A/H1N1pdm09 viruses, but that these fitness costs are modest and are not completely deleterious to viral replication and transmission.

In existing studies investigating the fitness of PA/I38T-substituted influenza viruses in animal models, variant influenza viruses of recombinant and patient-derived origins have been capable of robust viral replication in mice, hamsters and ferrets (Checkmahomed et al., 2019; Chesnokov et al., 2019; Imai et al., 2019; Jones et al., 2020). Ferrets represent the gold standard transmission model for evaluating the fitness of influenza viruses (Belser, Eckert, Tumpey, & Maines, 2016); of the available reports, patient-derived and recombinant I38T viruses have previously been shown to transmit from experimentally inoculated DFs to naïve recipients by direct (Jones et al., 2020) and airborne contact (Imai et al., 2019; Jones et al., 2020) routes. In our study, patient-derived A/H3N2 and A/H1N1pdm09 I38T-variants, when evaluated as pure viral populations, could sustain a chain of three sequential generations of direct contact transmission. However, competition experiments allowed us to determine more subtle differences in viral fitness, indicating that the treatment-emergent I38T substitution compromised direct contact transmissibility of the A/H1N1pdm09 isolate, whereas the same substitution appeared to more strongly attenuate within-host replication in the A/H3N2 isolate. Furthermore, a recombinant I38T-variant A/H1N1pdm09 virus demonstrated similar fitness characteristics in ferrets to the patient-derived A/H1N1pdm09 isolate, suggesting that transmission of the variant by respiratory droplets/aerosols was restricted to a greater extent than transmission by direct contact. Our study revealed that populations of I38T-variants are outcompeted by WT viruses in both A/H1N1pdm09 and A/H3N2; however, pure populations of I38T-variant viruses were still able to be transmitted in ferrets in accordance with previous studies (Imai et al., 2019; Jones et al., 2020).

Recent reports using in vivo competitive mixture models suggest that the fitness impact of I38T can vary based on the virus background. Checkmahomed and colleagues (Checkmahomed et al., 2019) reported that I38T-variants of recombinant viruses from 2009 (pandemic A/H1N1pdm09) and 2013 (seasonal A/H3N2) outcompeted cognate WT viruses within-host in mouse co-infections, whereas a separate study found a I38T-substituted A/H3N2 clinical isolate from 2017 to be mildly attenuated compared with WT in ferrets (Chesnokov et al., 2019). Furthermore, fitness observations from competition experiments in vitro have not always accurately predicted in vivo fitness characteristics (Checkmahomed et al., 2019). Such factors contribute to the difficulties associated with predicting the likely widespread emergence of a strain, even if the virus in question has equivalent or greater fitness than the respective WT virus being tested. A previous example of this has been observed following a competitive fitness ferret study to evaluate the fitness of an oseltamivir-resistant A/H1N1pdm09 virus with the H275Y NA substitution that caused a localised cluster of cases among untreated patients in Australia and Japan in 2011/12 (Butler et al., 2014; Hurt et al., 2012; Takashita et al., 2014). This study estimated that the oseltamivir-resistant variant virus had equivalent or even greater fitness relative to the cognate WT virus, thereby concluding that there was a significant risk that this virus would emerge and spread globally. Despite this finding in ferrets, no such dissemination of A/H1N1pdm09 H275Y oseltamivir-resistant viruses in the community has occurred in the intervening years (Lackenby et al., 2018).

Permissive mutations in the NA and haemagglutinin (HA) proteins were thought to have compensated for the intrinsic fitness costs of the H275Y mutation in former seasonal A/H1N1 viruses, thereby enabling their capacity to spread globally in 2008 (Baranovich et al., 2010; Bloom, Gong, & Baltimore, 2010; Takashita et al., 2015). This study utilised clinically derived and recombinant virus pairs, where the PA genes differed by the I38T substitution alone, demonstrating that this baloxavir-resistance substitution is detrimental to intrinsic viral fitness relative to WT. However, as I38T does not completely prohibit viral replication and transmission, the potential exists for additional permissive mutations to develop that recover any viral fitness loss associated with I38T, which may enable increasing prevalence in community circulation. While specific compensatory mutations in I38T-variants have not yet been identified, a previous report has highlighted an example of this process developing for influenza viruses with reduced susceptibility to favipiravir, another polymerase inhibitor drug. It was found that variant viruses harbouring a PB1 substitution conferring resistance to favipiravir caused a loss of viral fitness and polymerase activity, which was rescued by a compensatory substitution in another part of the polymerase complex to restore the fitness and polymerase function of the variant (Goldhill et al., 2018).

The error-prone properties of influenza virus replication mean that substitutions leading to resistance will inevitably occur. Amino acid substitutions that confer reduced susceptibility to both the adamantanes and NAIs are well described, and likewise treatment with baloxavir has been observed to select for variant viruses that have reduced baloxavir susceptibility in clinical trials. Resistant viruses may retain sufficient fitness to transmit from person to person, typically within households (Buchholz et al., 2010; Hayden et al., 1989). However, from a public health perspective it is those resistant viruses that demonstrate the potential to outcompete and displace circulating antiviral sensitive strains, as happened in 2007–2009 for the oseltamivir-resistant A/H1N1 virus, that are of significant concern. Our results in the ferret model provide evidence that the intrinsic fitness of I38T-substituted viruses with reduced baloxavir susceptibility is attenuated compared to baloxavir-sensitive WT strains. Indeed, despite an estimated 5.5 million doses of baloxavir supplied to Japanese medical institutions between October 2018 and January 2019 (Emi Takashita et al., 2019), there have been very few reports of potential human-to-human spread of a I38T-variant in Japanese households (E. Takashita et al., 2019; Emi Takashita et al., 2019). No clusters of sustained transmission indicating epidemiological spread of I38T-variants have been identified in any population or region to date and the number of baloxavir-resistant viruses detected in circulating viruses between September 2019 and January 2020 demonstrated that only a single E23K variant in 1355 A/H1N1pdm09 viruses, and a single I38M variant in 1012 A/H3N2 viruses, were detected in patients with no prior treatment (WHO, 2020). Thus, it appears that baloxavir usage has not resulted in a significant number of circulating viruses with reduced susceptibility to the drug. Nevertheless, our study suggests that the fitness impact of the I38T substitution in A/H1N1pdm09 and A/H3N2 influenza viruses is relatively modest, and the potential remains for the acquisition of permissive mutations to recover the fitness cost of baloxavir resistance.

## Materials and methods

### Cells and viruses

Canine kidney MDCK and MDCK-SIAT1 cells were cultured as previously described (*Supplementary File 1*) (Omoto et al., 2018; Uehara et al., 2019). The human nasal epithelial cells (MucilAir™) were obtained from Epithelix Sarl (Geneva, Switzerland) and the cells were maintained in MucilAir™ culture medium at 37 °C in a humidified 5% CO_2_ incubator.

Three A/H3N2 clinical isolate pairs were derived from human nasopharyngeal/pharyngeal swab samples from patients #344103, #253104, and #339111 from the CAPSTONE-1 phase 3 trial (otherwise-healthy patients with influenza; ClinicalTrials.gov identifier: NCT02954354) as previously described (Uehara et al., 2019). Two A/H1N1pdm09 clinical isolate pairs were derived from swabs from patients #2HB001 and #2PQ003 involved in a double-blind, multicentre, placebo-controlled phase 2 study in Japan (Japic CTI-153090) (*Figure 1 – figure supplement 3*). Pre-treatment samples contained viruses without an I38T substitution (WT) and post-treatment samples contained viruses with an I38T substitution; full-length HA, NA and PA sequences of clinical isolates from pre- and post-treatment samples were verified by Sanger sequencing (*Supplementary Table*). Clinical isolates were propagated and titrated in MDCK-SIAT1 cells. The selection criteria for clinical isolates used in the ferret study is summarised in *Figure 1 – figure supplement 2*.

For in vitro experiments, recombinant viruses based on A/WSN/33 (H1N1) and A/Victoria/3/75 (H3N2) were generated as previously described (Omoto et al., 2018). For in vivo experiments, a recombinant A/H1N1pdm09 virus pair of WT and I38T viruses were generated by reverse genetics using plasmids containing the gene segments from A/England/195/2009 as previously described (Frise et al., 2016). Recombinant viruses were propagated and titrated in MDCK cells.

### In vitro characterisation of clinical isolate pairs

The baloxavir sensitivity of clinical isolate pairs (WT and I38T) was assessed by plaque reduction assay in MDCK-SIAT1 cells as previously described (Uehara et al., 2019). Baloxavir acid was synthesised by Shionogi & Co., Ltd. (Osaka, Japan). The effective concentration of baloxavir achieving 50% inhibition of plaque formation (EC_50_) was calculated for each virus based on plaque numbers. The mean (± SD) EC_50_ was determined from three independent experiments performed in duplicate. Data for A/H3N2 viruses (#344103, #253104, and #339111) were previously reported by Uehara et al. (Uehara et al., 2019).

The replication of clinical isolate virus pairs was compared in vitro using primary human airway epithelial MucilAir™ cells. Confluent monolayers of MucilAir™ cells in transwell 24-well plates (5.0×10^5^ cells/well) were infected with either WT or I38T virus at a multiplicity of infection (MOI) of 0.001 for both A/H1N1pdm09 and A/H3N2 viruses. The culture supernatants were collected at 0, 24, 48 and 72 hours after inoculation and the viral titre was determined by TCID_50_ measured in MDCK-SIAT1 cells.

Competitive fitness experiments were conducted in MucilAir™ cells using A/H3N2 and A/H1N1pdm09 clinical isolate pairs. The WT virus and the virus harbouring the I38T substitution were diluted to equivalent infectious viral titres calculated by TCID_50_ in MDCK-SIAT1 cells and mixed at 50:50 or 20:80 volumetric ratios. Confluent monolayers of MucilAir™ cells in transwell-24 plates were infected at an MOI of 0.002. Cell culture supernatant was collected 48 hours post infection and viral titres determined by TCID_50_. The virus supernatant were serially passaged three times in duplicate cultures at an MOI of 0.001, and the proportion of viruses harbouring I38X substitutions were evaluated by deep sequencing as previously described (Uehara et al., 2019). The PA coding region (nucleotides 1–466; amino acids 1–155 for A/H3N2 viruses and nucleotides 1–419; amino acids 1–139 for A/H1N1pdm09 viruses) were amplified from total RNA extracted from each culture supernatant (MagNA Pure 96 system; Roche Life Science, Superscript III reverse transcriptase; Invitrogen, HotStar Taq DNA polymerase; Qiagen, Venlo, The Netherlands). Fragmented and barcoded DNA libraries were generated (Nextera XT Library Prep Kit; Illumina) and sequenced using one flow cell on an Illumina MiSeq instrument. All reads were mapped to the PA gene of the reference strain A/Texas/50/2012 (H3N2) or A/California/07/2009 (H1N1) using the CLC Genomics Workbench Version 10.0.1. The average coverage was more than 10,000 and the percentage of reads with a quality score greater than 30 was more than 69%. A threshold frequency of >1% was adopted for calling variants, and the calculated risk of false positive variants was 0.21% for A/H3N2 and 0% for A/H1N1pdm09 strains, respectively, which were determined by sequencing duplicate influenza A virus stock samples.

### Ferret studies

#### Melbourne experiments

The replication and transmission of clinical isolate virus pairs was evaluated in vivo using a competitive fitness experiment design in ferrets (*Figure 3 – figure supplement 1*). Outbred male and female ferrets (*Mustela putorius furo*) >12 weeks old (independent vendors), weighing 600–1800 g and seronegative against recently circulating influenza viruses were used.

Virus stocks for ferret inoculation were diluted to equivalent viral titres in PBS (10^5^ TCID_50_/500 μL for the A/H3N2 viruses and 10^4^ TCID_50_/500 μL for the A/H1N1pdm09 viruses) and the WT and I38T-variant pairs were then mixed at various volumetric ratios: 100% WT : 0% I38T, 80% WT : 20% I38T, 50% WT : 50% I38T (×2 groups), 20% WT : 80% I38T, and 0% WT : 100% I38T; n = 4 ferrets per transmission chain, total n = 24 ferrets per experiment). Within each transmission chain, DFs were anaesthetised by intramuscular injection of a 1:1 (v/v) xylazine/ketamine (10 mg/kg, Troy Laboratories) mixture, and inoculated with pure or mixed virus suspension (250 μL per nostril) by the intranasal route. The first naïve recipient (RF1) was exposed to the DF 1 DPI of virus. Ferrets were nasal washed daily under xylazine sedation (5 mg/kg, Troy Laboratories) and confirmed to be influenza positive based on a real-time reverse-transcriptase polymerase chain reaction (RT-PCR) cycle threshold (C_t_) of <35. Following detection of virus in RF1, it was cohoused 1:1 with a subsequent naïve recipient (RF2) the next day. Similarly, once influenza was detected in RF2 it was cohoused with RF3, and the transmission chain was terminated.

Daily nasal washes were collected under xylazine sedation (0.5 mg/kg) by instilling 1 mL PBS (1% w/v BSA) into the nostrils for up to 10 consecutive days following inoculation/exposure. Nasal wash samples were stored at –80 °C until determination of infectious virus titres by in MDCK-SIAT1 cells as previously described (Oh, Barr, & Hurt, 2015). The lower limit of quantitation (LLOQ) was 2.0 log_10_ TCID_50_/mL. The experiment was terminated at 16 DPI, regardless of transmission status.

#### London experiments

Female ferrets (20–24 weeks old) weighing 750–1000 g were used (independent vendors). After acclimatisation, sera were obtained and tested by HI assay for antibodies against A/England/195/2009/H1N1pdm09. All ferrets were confirmed to be seronegative against the NP protein of influenza A virus by using an ID Screen Influenza virus A antibody competition enzyme-linked immunosorbent assay kit (ID.vet) at the start of the experiments.

DFs (n = 4) were anaesthetised with ketamine (22 mg/kg) and xylazine (0.9 mg/kg) for intranasal inoculation with 10^4^ PFU of the 50% WT : 50% I38T mixture of the reverse genetics A/England/195/1009 virus pair diluted in PBS (100 μL per nostril). The volumetric ratio of virus inoculation was determined based on infectious virus titre (PFU/mL). At 24 hours post inoculation, a direct contact recipient was exposed by cohousing with the donor in the same cage, while simultaneously an indirect contact recipient was placed in an adjacent cage (minimum 6 cm apart) which only allowed respiratory droplet/aerosol exposure from the donor cage (*Figure 6 – figure supplement 1*). Exposure of recipient animals to donor animals (1:1:1 ratio) was continued until 3 DPI (48-hour exposure period).

All animals were nasal washed daily, while conscious, by instilling 2 mL PBS into the nostrils. The nasal wash sample was used for virus titration by plaque assay in MDCK cells. The LLOQ in the plaque assays was 10 PFU/mL.

### Quantitative real-time RT-PCR (qRT-PCR)

For the Melbourne experiments, viral RNA was extracted from 200 μL nasal wash samples using the NucleoMag VET isolation kit (Macherey Nagel) on the KingFisher Flex (ThermoFisher Scientific) platform according to manufacturer’s instructions. Primer/probe sets were obtained from CDC Influenza Division: Influenza virus M gene copy number per μL RNA was determined by qRT-PCR using the SensiFAST Probe Lo-ROX One-Step qRT-PCR System Kit (Bioline) on the ABI 7500 Real Time PCR System (Applied Biosystems) under the following conditions: 45 °C for 10 min, one cycle; 95 °C for 2 min, one cycle; 95 °C for 5 s then 60 °C for 30 s, 40 cycles. Influenza A genomic copies were quantitated by the standard curve method using influenza A RNA samples of known copy number generated in-house. The M gene-targeted primers/TaqMan probe for the detection of influenza A viruses are sourced from the CDC Influenza Division (Atlanta, USA): forward primer (5’-ACCRATCCTGTCACCTCTGAC-3’), reverse primer (5’-GGGCATTYTGGACAAAKCGTCTACG-3’) and TaqMan probe (6FAM-TGCAGTCCTCGCTCACTGGGCACG-BHQ1). Results were analysed by 7500 Fast System SDS software v1.5.1. The LLOQ was 100 M gene copies/μL RNA, based on the first RNA standard to yield C_t_ <35.

For the London experiments, viral RNA was extracted from 140 μL nasal wash samples using the QIAamp viral RNA mini kit (Qiagen) according to manufacturer’s instructions. Real-time RT-PCR was performed using 7500 Real Time PCR system (ABI) in 20 μL reactions using AgPath-ID™ One-Step RT-PCR Reagents 10 µl RT-PCR buffer (2X) (Thermo Fisher), 4 μL of RNA, 0.8 µl forward (5’-GACCRATCCTGTCACCTCTGA-3’) and reverse primers (5’-AGGGCATTYTGGACAAAKCGTCTA-3’) and 0.4 µl probe (5’-FAM-TCGAGTCCTCGCTCACTGGGCACG-BHQ1-3’). The following conditions were used: 45 °C for 10 min, one cycle; 95 °C for 10 min, one cycle; 95 °C for 15 s then 60 °C for 1 min, 40 cycles. For each sample, the C_t_ value for the target M gene was determined, and absolute M gene copy numbers were calculated based on the standard curve method. The LLOQ was 153 M gene copies/μL RNA, based on the results from the samples of uninfected ferrets (mean + 2*SD).

### Quantitation of I38T-variant population for ferret studies

For the Melbourne experiments, detection of the I38T-variant in a virus sample was performed by pyrosequencing as previously described (Koszalka, Farrukee, Mifsud, Vijaykrishna, & Hurt, 2020). The MyTaq One-Step RT-PCR Kit (Bioline) was used to generate and amplify cDNA from viral RNA isolated from nasal wash samples (as above). Samples containing fewer than 200 M gene copies/μL RNA were excluded from analysis. RT-PCR product was processed in the PyroMark Vacuum Workstation according to manufacturer’s protocol, and pyrosequencing was performed using the PyroMark ID System (Biotage). Primers were designed to investigate SNPs at position 38 of the PA gene of influenza A/H3N2 and A/H1N1pdm09 (*Supplementary File 1*). Analysis was performed using PyroMarkQ96 ID Software to quantify the percentage of WT and variant in a mixed virus population. The error associated with estimating pure populations of WT or I38T virus was such that values <5% or >95% could not be accurately quantified – therefore, values <5% or >95% were considered to be pure WT or I38T, respectively (Koszalka et al., 2020).

For the London experiments, whole-genome NGS was performed using a pipeline at Public Health England (PHE). Viral RNA was extracted from viral lysate using easyMAG (bioMérieux). One-step RT-PCR was performed using SuperScript III (Invitrogen), Platinum Taq HiFi polymerase (Thermo Fisher), and influenza-specific primers (Zhou et al., 2009). The entire ORF of the PA segment was amplified in two overlapping fragments using two pairs of primers (*Supplementary File 1*). Samples were prepared using the Nextera XT library preparation kit (Illumina) and sequenced on an Illumina MiSeq generating a 150-bp paired-end reads. Reads were mapped with BWA (version 0.7.5) and converted to BAM files using SAMTools (version 1.1.2). Variants were called using QuasiBAM, an in-house script at PHE. Analysis of prepared minority variant mixture controls has shown that a cut-off set at 5% coupled with a minimum depth of 2000 were reliable parameters to quantify the I38T-variant with 99% confidence (data not shown).

We considered the I38T-variant proportion measured at experiment endpoint as dependent on both within-host and between-host components of viral fitness. Within-host fitness was regarded as the observed trend in the proportion of %I38T changing in each individual ferret over time. Between-host fitness was regarded as the observed differences in the measured proportion of I38T-variant in the RF compared to the proportion measured in the DF on the previous day for each transmission event.

### Statistical analysis

For in vitro experiments, comparisons of virus titres between WT and I38T viruses at each time-point were conducted using Welch’s *t*-test by using the statistical analysis software, SAS version 9.2 at the 0.05 significance level using two-sided tests. No adjustment for multiple testing was performed for exploratory analysis.

For the Melbourne in vivo experiments, 2-way ANOVA followed by Bonferroni’s post-tests (Prism 5) were performed on daily virus titre data from pure WT- and I38T-group ferrets which were infected by direct contact transmission (n = 3 per group). Data from DF were not included in this analysis as the replication kinetics of experimental infection by direct inoculation of virus differs from the surrogate for ‘natural’ infection that is achieved by direct contact. AUC analysis was performed, and AUC values for WT and I38T groups were compared by two-tailed Mann-Whitney U test (Prism 5). p≤0.05 was considered statistically significant.

### Ethics

The Melbourne ferret experiments were conducted with approval (AEC#1714278) from the University of Melbourne Biochemistry & Molecular Biology, Dental Science, Medicine, Microbiology & Immunology, and Surgery Animal Ethics Committee, in accordance with the NHMRC Australian code of practice for the care and use of animals for scientific purposes (8th edition). For the London ferret experiments, all work performed was approved by the local genetic manipulation (GM) safety committee of Imperial College London, St. Mary’s Campus (centre number GM77), and the Health and Safety Executive of the United Kingdom.

## Acknowledgements

The Melbourne WHO Collaborating Centre for Reference and Research on Influenza is supported by the Australian Government Department of Health. Editorial support was provided by John Bett, PhD, of Gardiner-Caldwell Communications (GCC), Macclesfield, UK. This assistance was funded by F. Hoffmann-La Roche Ltd.

## Additional files

### Supplementary files

- Supplementary file 1. Supplemental methods and figures

## Supplementary file 1

### Supplementary methods

- Cell culture
  - MDCK-SIAT1 cells were obtained from the European Collection of Cell Cultures and were grown in Dulbecco’s modified Eagle’s medium (DMEM) supplemented with 2 mM L-glutamine solution (Nacalai Tesque, Inc.), 1 µg/mL Geneticin™ selective antibiotic (Thermo Fisher Scientific, Inc.), 10% fetal bovine serum (FBS) (Sigma-Aldrich Co., Ltd.), and penicillin-streptomycin (Thermo Fisher Scientific, Inc.).
- Virus titration
  - The virus titre of clinical isolates was determined by the 50% tissue culture infectious dose (TCID_50_) method in MDCK-SIAT1 cells in DMEM containing 0.2% bovine serum albumin (BSA) (Sigma-Aldrich Co., Ltd.), 25 mM HEPES (Thermo Fisher Scientific, Inc.), 2 mM L-glutamine solution, 1 µg/mL trypsin from bovine pancreas-TPCK treated (TPCK trypsin) solution (Sigma-Aldrich Co., Ltd.), and penicillin-streptomycin.
- In vitro competitive fitness of rg virus pairs:
  - Competitive fitness experiments were conducted in MDCK cells using 50:50 mixtures of WT/I38X virus pairs prepared volumetrically based on infectious viral titres (TCID_50_). The presence of I38X variant in cell culture supernatant was determined by Sanger sequencing.

Melbourne methods:

- Cell culture
  - MDCK-SIAT1 cells were obtained from ATCC. Cultures were grown in Dulbecco’s modified Eagle’s medium (DMEM) supplemented with 10% (v/v) foetal bovine serum, 2 mM GlutaMAX, 0.05% sodium bicarbonate, 100 μM MEM non-essential amino acid, 20 mM HEPES, 500mg Geneticin and 50,000 U penicillin-streptomycin.
- Virus titration
  - The virus titre of clinical isolates was determined by the TCID_50_ method in MDCK-SIAT1 cells in DMEM containing 2 mM GlutaMAX, 0.05% sodium bicarbonate, 100 μM MEM non-essential amino acid, 20 mM HEPES, 100,000 U penicillin-streptomycin, 1000 μg Amphotericin B and 4 μg/mL of TPCK-treated trypsin.
- Ferret handling procedures
  - Identification/temperature-monitoring chips (LifeChip, Bio-Thermo) were implanted subcutaneously on the dorsal region of each ferret. Animals were monitored at least once daily for temperature and weight changes, had *ad libitum* access to pellet feed (Eukanuba) and water and were provided with supplementary wet food (Hill’s Pet Nutrition, Australia) as ethically required (below 90% baseline weight). Nasal wash samples were collected using 1 mL of PBS containing 1% w/v BSA. At the end of each experiment, ferrets were anaesthetised before sacrifice by intracardiac injection of sodium pentobarbitone (≤1,000 mg/kg, Troy Laboratories).
- Pyrosequencing primers (Melbourne)
  - A/H3N2: Forward (Biotin-5’-TTGTCGAACTTGCAGAAAAGGC-3’), reverse (5’-GCCATTGTTCTGTCTCTCCCCT-3’) and pyrosequencing (5’-CATACCTCCAAGTGAGTGCA-3’, reverse orientation)
  - A/H1N1pmd09: Forward (Biotin-5’-CAATCCAATGATCGTCGAGC-3’), reverse (5’-GGTGCTTCAATAGTGCATTTGG-3’) and pyrosequencing (5’-CAAACTTCCAAATGTGTGCA-3’, reverse orientation)

London methods:

- Ferret handling
  - Body weight was measured daily, and strict procedures were followed to prevent aberrant cross-contamination between animals. Sentinel animals were handled before inoculated animals, and work surfaces and handlers’ gloves were decontaminated between animals.
- Cell culture:
  - MDCK cells were maintained in Dulbecco’s modified Eagle’s medium (DMEM; Gibco, Invitrogen) supplemented with 10% FBS and 1% penicillin/streptomycin (Sigma-Aldrich) and 1% non-essential amino acid (Gibco, Invitrogen).
- Plaque assays:
  - 100% confluent MDCK cell monolayers were inoculated with 100 μL of serially diluted samples and overlaid with 0.6% agarose (Oxoid) in supplemented DMEM with 1 μg trypsin (Worthington) mL−1 and incubated at 37 °C for 3 days.
- Next-generation sequencing primer sets:
  - Full genome sequencing as described previously (Zhou et al., 2009)
  - PA ORF amplified in two overlapping fragments PA-1 and PA-2. PA-1 forward (5’-AGCAAAAGCAGGTACTGATCCAA-3’), PA-1 reverse (5’-CTTGAATCAGTCAATTCACATGCC-3’), PA-2 forward (5’-ATGCAATCAAATGCATGAAGA-3’), PA-2 reverse 5’-ATATGGTCTCGTATTAGTAGAAACAAGGTACTT-3’)

**Supplementary Table.** Amino acid and nucleotide difference in A/H1N1pdm09 and A/H3N2 viruses isolated from baloxavir-treated patients.

|  | Segment | A/H1N1pdm09 |  | A/H3N2 |  |  |
| --- | --- | --- | --- | --- | --- | --- |
|  |  | 2HB001 | 2PQ003 | 344103 | 339111 | 253104 |
| Amino acid | HA | -* | - | - | - | - |
|  | NA | - | - | - | - | - |
|  | PA | I38T | I38T | I38T | I38T | I38T |
| **Nucleotide** | HA | - | - | t885c | - | - |
|  | NA | g1245a | - | - | - | a732g |
|  | PA | t113c | t113c | t113c | t113c<br>g1698a | t113c |

\*No mutation was detected compared with the cognate wild type virus in amino acid or nucleotide. HA, haemagglutinin NA, neuraminidase; PA, polymerase acidic

## Supplementary figures

**Figure 1 – figure supplement 1.**
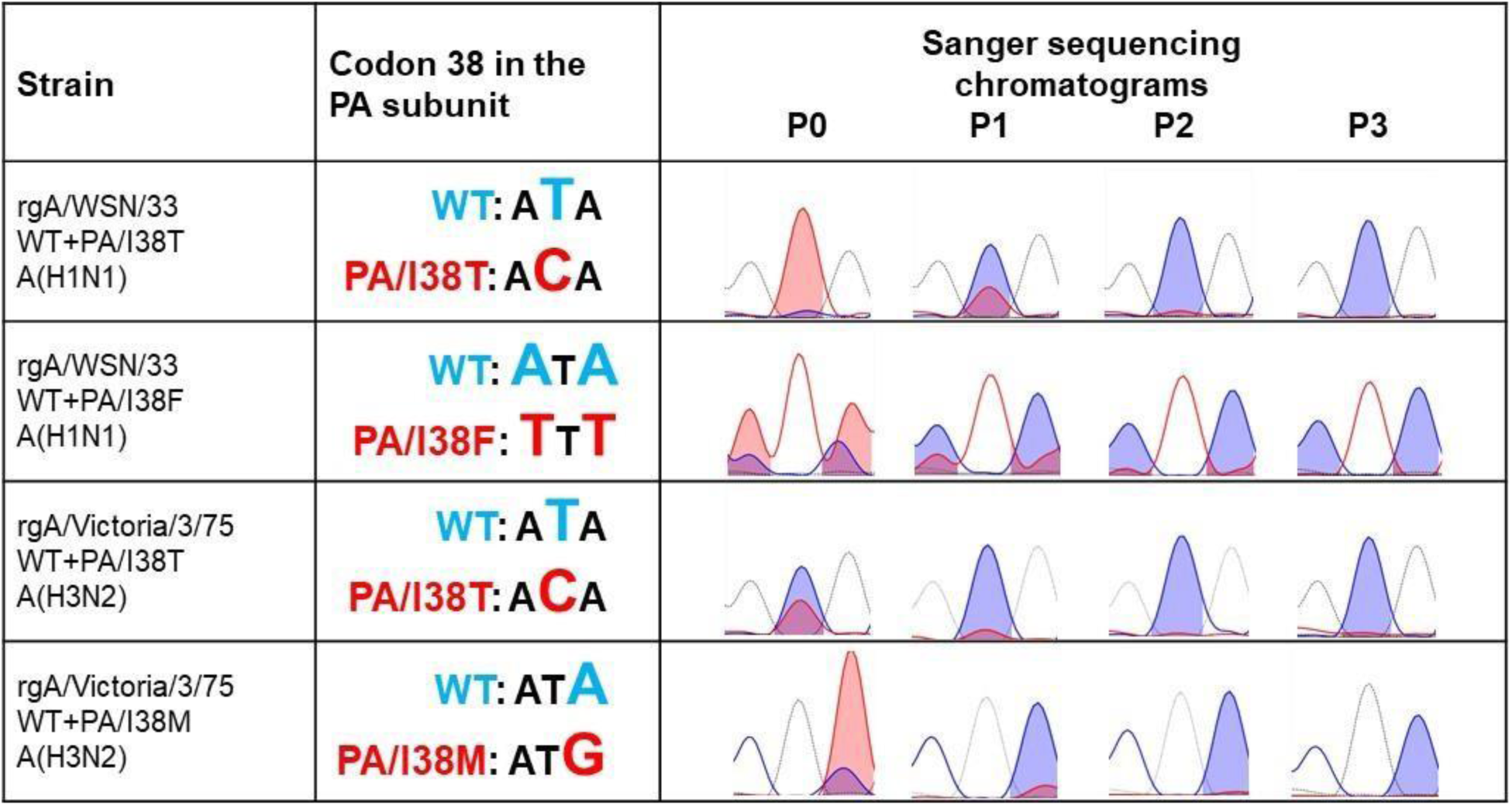
Competitive replicative capacity of influenza reverse genetics-derived viruses with PA/I38 substitutions. Competitive replicative capacity of reverse genetics-derived influenza viruses with I38 substitutions. MDCK cells were co-infected with reverse genetics-derived WT and I38-substitutited viruses at 50:50 ratio based on viral titres at an MOI of 0.001 each. At 48 hours post infection, the culture supernatant was collected and serially passaged three times. The virus passage experiments were conducted in duplicate as lineage 1 and lineage 2. Each culture supernatant was subjected to Sanger sequencing to analyse the change of amino acid at 38 in PA subunit. Sanger sequence chromatograms of amino acid at position 38 in the PA subunit were analysed using BioPython, and blue chromatograph represents WT and red shows /I38-substituted viruses. Representative sequencing chromatographs of lineage 1 were shown because the similar results were obtained from both lineages. MOI, multiplicity of infection; PA, polymerase acid; P0–P3, passage 0 to passage 3; WT, wild type.

**Figure 1 – figure supplement 2.**
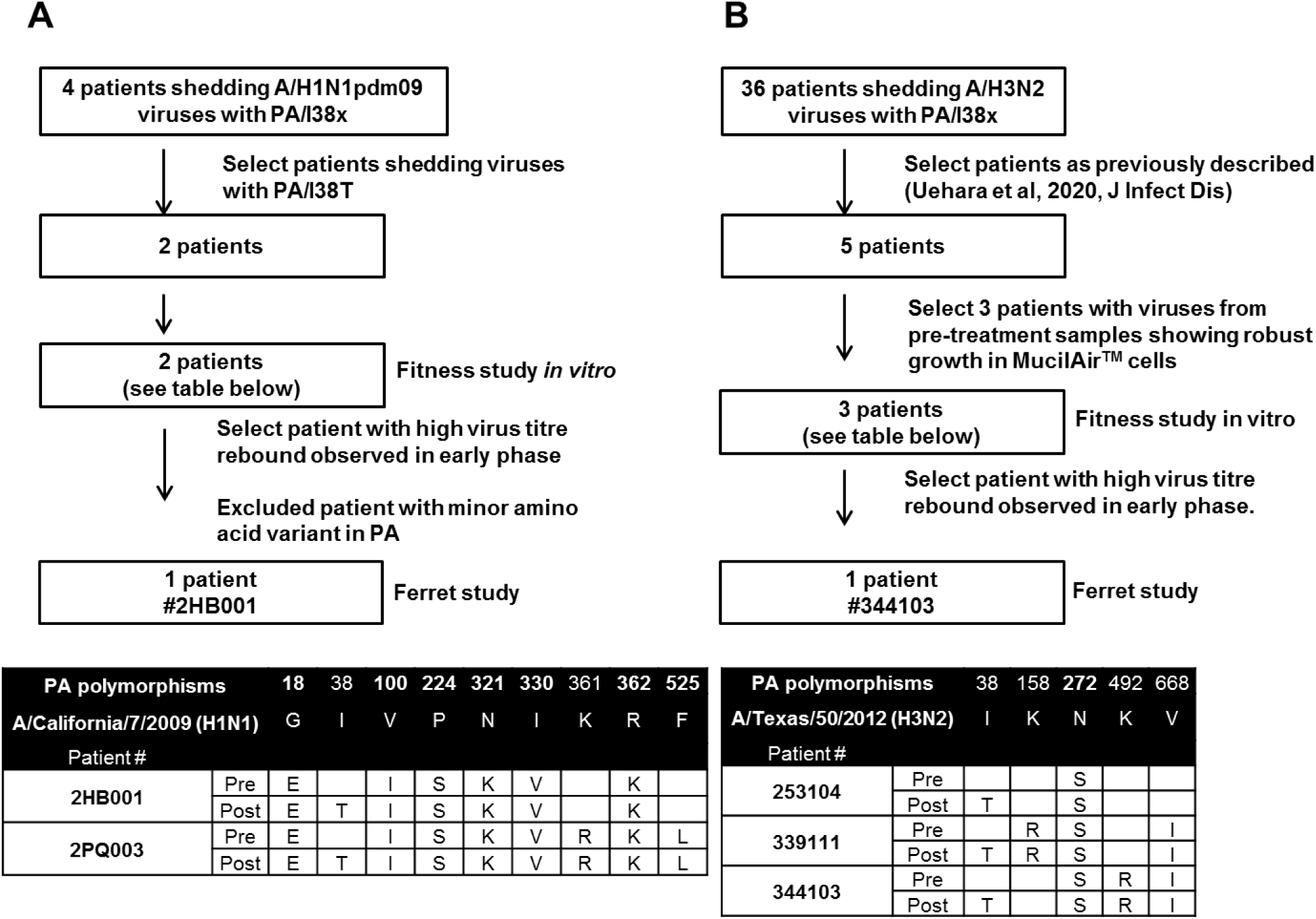
Sample selection for fitness study in vitro and ferret transmission study. Sample selection process for in vitro fitness and ferret transmission studies of I38-substituted viruses propagated from swab samples. Flow diagram of sample selection for fitness study in vitro and ferret transmission study for (A) influenza A/H1N1pdm09 and (B) influenza A/H3N2 viruses. Polymorphic amino acid substitutions in PA protein aligned with (A) influenza A/California/7/2009 (A/H1N1) and (B) influenza A/Texas/50/2012 (A/H3N2) are also shown. PA, polymerase acidic.

**Figure 1 – figure supplement 3.**
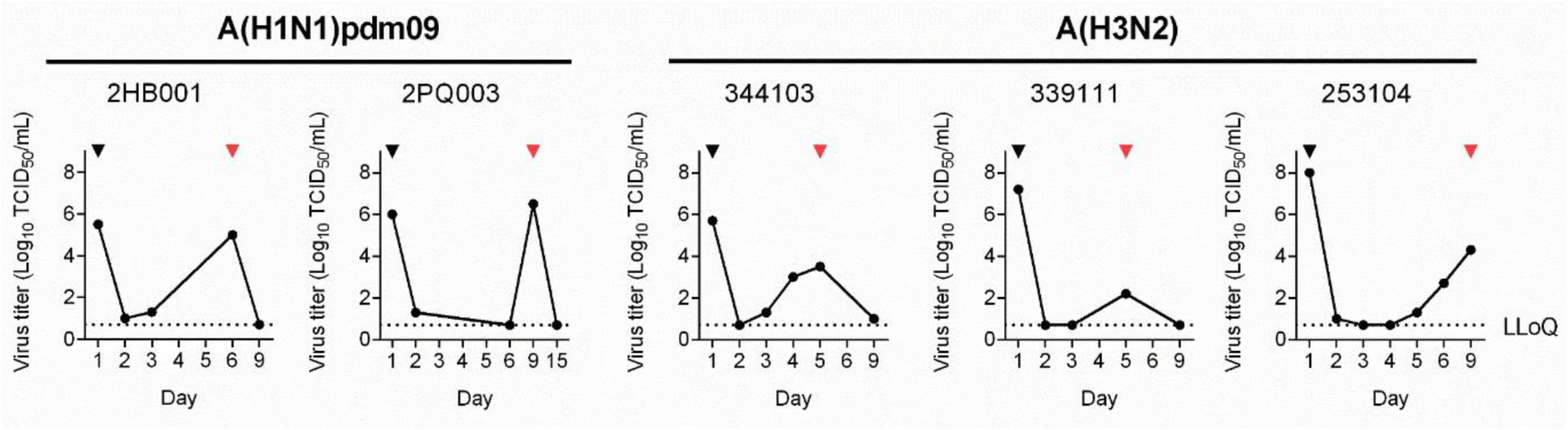
Influenza virus titre (log10 TCID50/mL) in individual patients with PA/I38T-substituted viruses. Influenza virus titre (log_10_ TCID_50_/mL) in individual patients with I38T-substituted viruses. Time course of influenza virus titre of individual baloxavir-treated patients with I38X-substituted viruses, determined by TCID_50_ assay (values below LLOQ were set at 0.7 log_10_ TCID_50_/mL). The black and red arrowheads indicate the sampling time-points for WT viruses (prebaloxavir treatment) and I38T-substituted viruses (post-baloxavir treatment), respectively. Black dotted line = LLOQ at 0.7 log_10_ TCID_50_. LLOQ, lower limit of quantitation; TCID_50_, 50% tissue culture infectious dose.

**Figure 2 – figure supplement 1.**
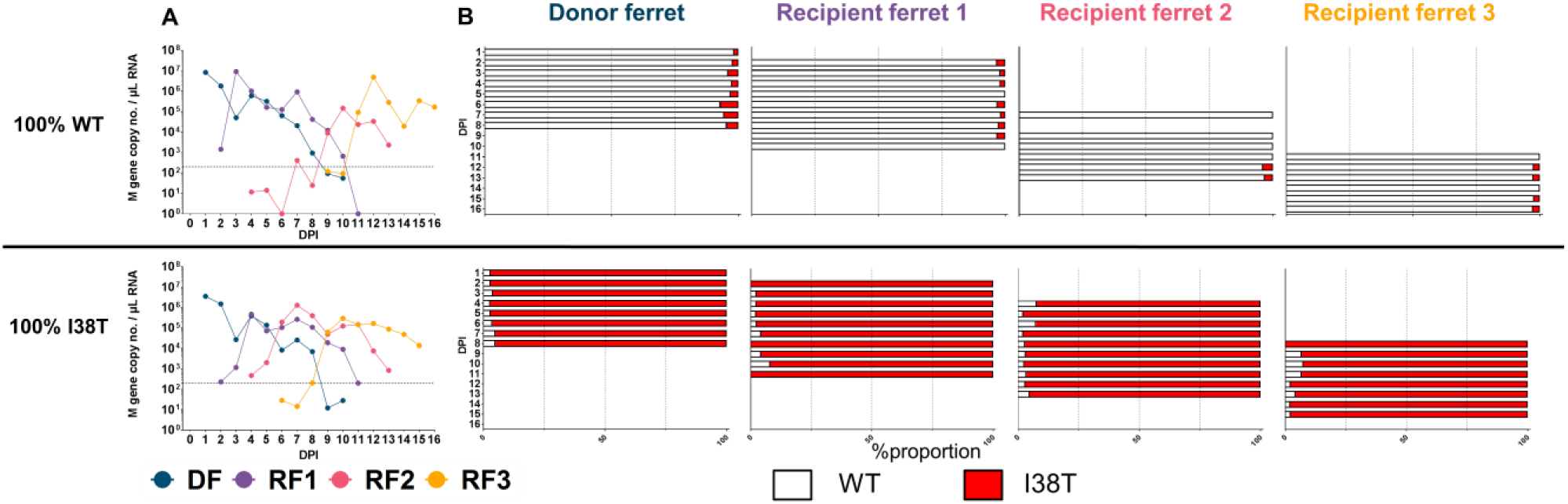
Pyrosequencing of ferret nasal washes from WT or I38T-variant A/H3N2 pure population groups. (**A**) Viral RNA and (**B**) pyrosequencing of ferret nasal washes from A/H3N2 WT or I38T pure population groups. DPI, days post inoculation; WT, wild type.

**Figure 2 – figure supplement 2.**
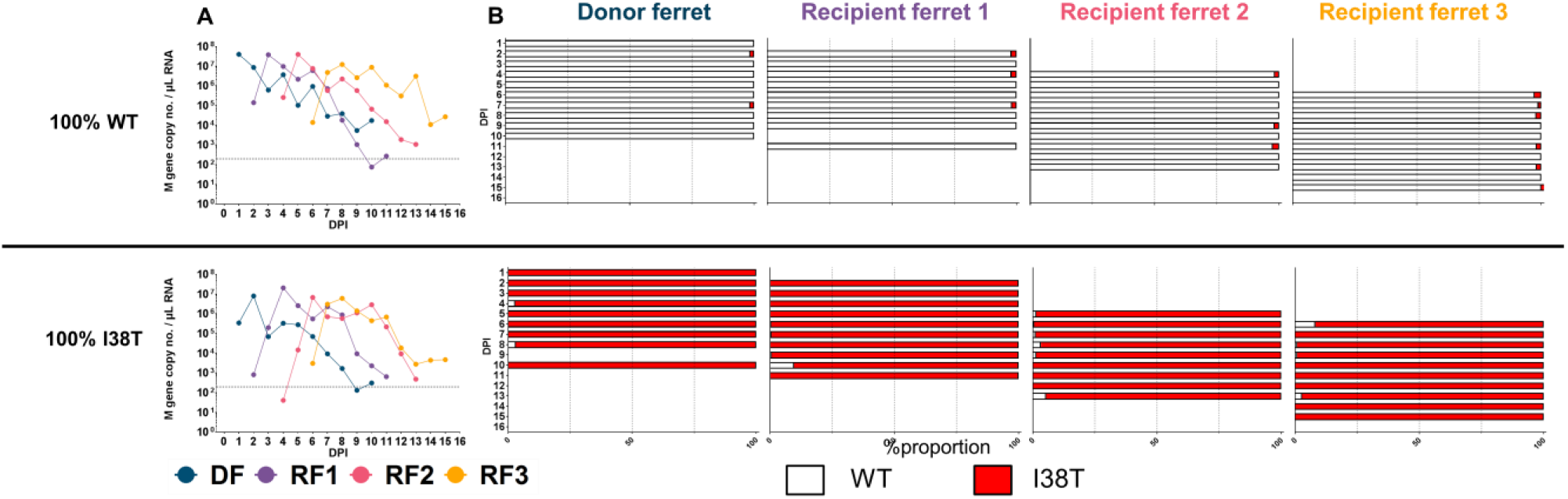
Pyrosequencing of ferret nasal washes from WT or I38T-variant A/H1N1pdm09 pure population groups. (A) Viral RNA and (B) pyrosequencing of ferret nasal washes from A/H1N1pdm09 WT or I38T pure population groups. DPI, days post inoculation; WT, wild type.

**Figure 3 – figure supplement 1.**
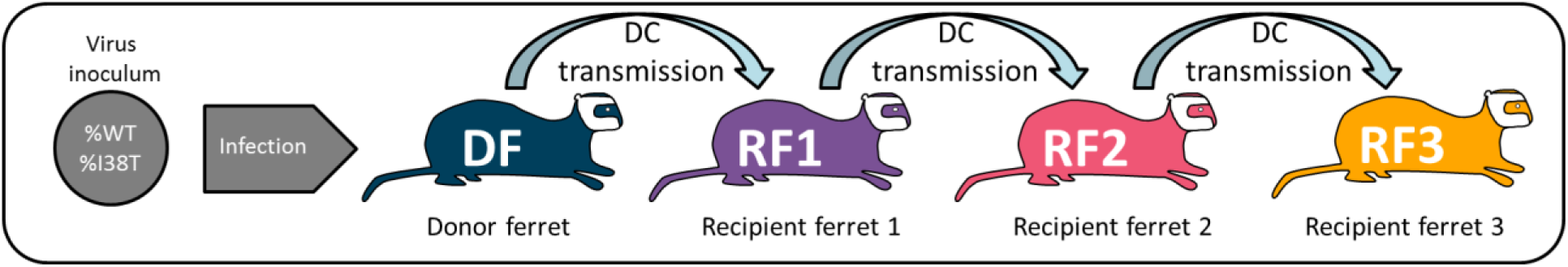
Ferret experiment designs. Ferret transmission experiment design schematics for Melbourne clinical isolate studies. DC, direct contact; DF, donor ferret; RF, recipient ferret; WT, wild type.

**Figure 6 – figure supplement 1.**
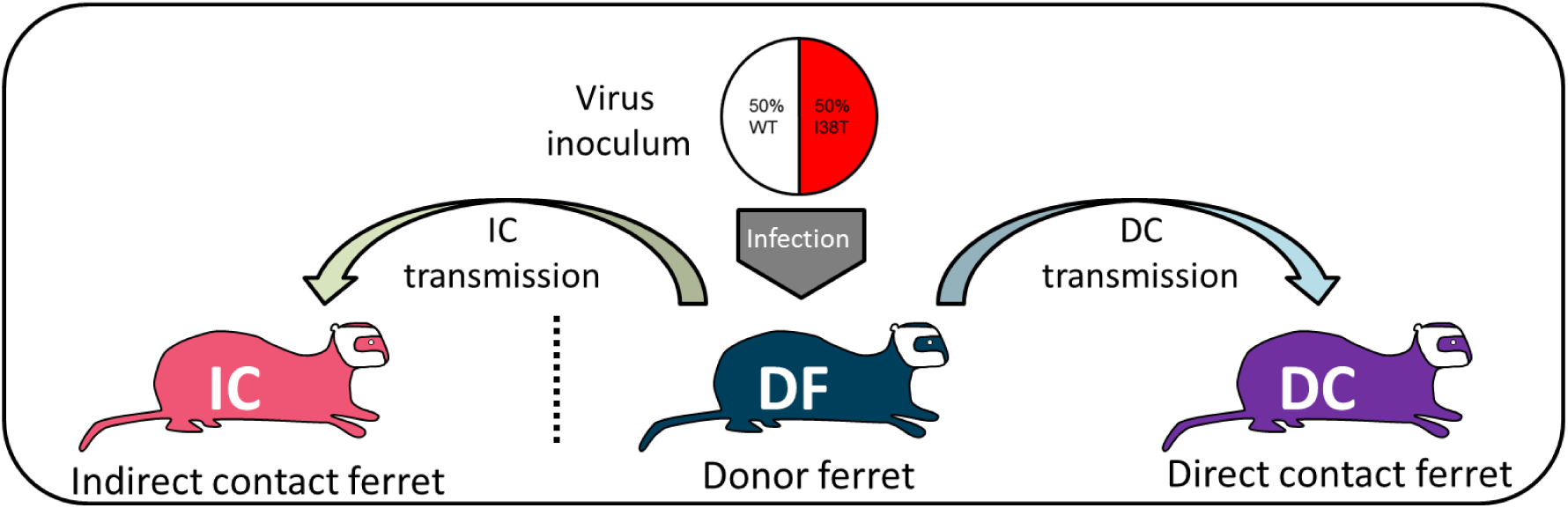
Ferret experiment designs. Ferret transmission experiment design schematics the London recombinant virus study. DC, direct contact; DF, donor ferret; IC, indirect contact; WT, wild type.

## References

Abed, Y., Goyette, N., & Boivin, G. (2005). Generation and Characterization of Recombinant Influenza A (H1N1) Viruses Harboring Amantadine Resistance Mutations. Antimicrobial Agents and Chemotherapy, 49(2), 556–559. doi: 10.1128/aac.49.2.556-559.2005

Baker, J., Block, S. L., Matharu, B., Burleigh Macutkiewicz, L., Wildum, S., Dimonaco, S., … Piedra, P. A. (2020). Baloxavir Marboxil Single-dose Treatment in Influenza-infected Children: A Randomized, Double-blind, Active Controlled Phase 3 Safety and Efficacy Trial (miniSTONE-2). Pediatr Infect Dis J. doi: 10.1097/INF.0000000000002747

Baranovich, T., Saito, R., Suzuki, Y., Zaraket, H., Dapat, C., Caperig-Dapat, I., … Suzuki, H. (2010). Emergence of H274Y oseltamivir-resistant A(H1N1) influenza viruses in Japan during the 2008-2009 season. J Clin Virol, 47(1), 23–28. doi: 10.1016/j.jcv.2009.11.003

Belser, J. A., Eckert, A. M., Tumpey, T. M., & Maines, T. R. (2016). Complexities in Ferret Influenza Virus Pathogenesis and Transmission Models. Microbiology and molecular biology reviews : MMBR, 80(3), 733–744. doi: 10.1128/MMBR.00022-16

Bloom, J. D., Gong, L. I., & Baltimore, D. (2010). Permissive secondary mutations enable the evolution of influenza oseltamivir resistance. Science (New York, N.Y.), 328(5983), 1272–1275. doi: 10.1126/science.1187816

Buchholz, U., Brockmann, S., Duwe, S., Schweiger, B., an der Heiden, M., Reinhardt, B., & Buda, S. (2010). Household transmissibility and other characteristics of seasonal oseltamivir-resistant influenza A(H1N1) viruses, Germany, 2007-8. Euro Surveill, 15(6).

Butler, J., Hooper, K. A., Petrie, S., Lee, R., Maurer-Stroh, S., Reh, L., … Hurt, A. C. (2014). Estimating the Fitness Advantage Conferred by Permissive Neuraminidase Mutations in Recent Oseltamivir-Resistant A(H1N1)pdm09 Influenza Viruses. PLOS Pathogens, 10(4), e1004065. doi: 10.1371/journal.ppat.1004065

Checkmahomed, L., M’hamdi, Z., Carbonneau, J., Venable, M.-C., Baz, M., Abed, Y., & Boivin, G. (2019). Impact of the baloxavir-resistant polymerase acid (PA) I38T substitution on the fitness of contemporary influenza A(H1N1)pdm09 and A(H3N2) strains. The Journal of Infectious Diseases. doi: 10.1093/infdis/jiz418

Chesnokov, A., Patel, M. C., Mishin, V. P., De La Cruz, J. A., Lollis, L., Nguyen, H. T., … Gubareva, L. V. (2019). Replicative Fitness of Seasonal Influenza A Viruses With Decreased Susceptibility to Baloxavir. The Journal of Infectious Diseases, 221(3), 367–371. doi: 10.1093/infdis/jiz472

Dharan, N. J., Gubareva, L. V., Meyer, J. J., Okomo-Adhiambo, M., McClinton, R. C., Marshall, S. A., … Oseltamivir-Resistance Working Group, f. t. (2009). Infections With Oseltamivir-Resistant Influenza A(H1N1) Virus in the United States. JAMA, 301(10), 1034–1041. doi: 10.1001/jama.2009.294

Dias, A., Bouvier, D., Crepin, T., McCarthy, A. A., Hart, D. J., Baudin, F., … Ruigrok, R. W. (2009). The cap-snatching endonuclease of influenza virus polymerase resides in the PA subunit. Nature, 458(7240), 914–918. doi: 10.1038/nature07745

Dong, G., Peng, C., Luo, J., Wang, C., Han, L., Wu, B., … He, H. (2015). Adamantane-Resistant Influenza A Viruses in the World (1902–2013): Frequency and Distribution of M2 Gene Mutations. PLOS ONE, 10(3), e0119115. doi: 10.1371/journal.pone.0119115

Duan, S., Boltz, D. A., Seiler, P., Li, J., Bragstad, K., Nielsen, L. P., … Govorkova, E. A. (2010). Oseltamivir–Resistant Pandemic H1N1/2009 Influenza Virus Possesses Lower Transmissibility and Fitness in Ferrets. PLOS Pathogens, 6(7), e1001022. doi: 10.1371/journal.ppat.1001022

Frise, R., Bradley, K., van Doremalen, N., Galiano, M., Elderfield, R. A., Stilwell, P., … Barclay, W. S. (2016). Contact transmission of influenza virus between ferrets imposes a looser bottleneck than respiratory droplet transmission allowing propagation of antiviral resistance. Sci Rep, 6, 29793. doi: 10.1038/srep29793

Goldhill, D. H., te Velthuis, A. J. W., Fletcher, R. A., Langat, P., Zambon, M., Lackenby, A., & Barclay, W. S. (2018). The mechanism of resistance to favipiravir in influenza. Proceedings of the National Academy of Sciences, 115(45), 11613–11618. doi: 10.1073/pnas.1811345115

Hayden, F. G., Belshe, R. B., Clover, R. D., Hay, A. J., Oakes, M. G., & Soo, W. (1989). Emergence and apparent transmission of rimantadine-resistant influenza A virus in families. N Engl J Med, 321(25), 1696–1702. doi: 10.1056/nejm198912213212502

Hayden, F. G., Sugaya, N., Hirotsu, N., Lee, N., de Jong, M. D., Hurt, A. C., … Watanabe, A. (2018). Baloxavir Marboxil for Uncomplicated Influenza in Adults and Adolescents. New England Journal of Medicine, 379(10), 913–923. doi: 10.1056/NEJMoa1716197

Heneghan, C. J., Onakpoya, I., Jones, M. A., Doshi, P., Del Mar, C. B., Hama, R., … Jefferson, T. (2016). Neuraminidase inhibitors for influenza: a systematic review and meta-analysis of regulatory and mortality data. Health Technol Assess, 20(42), 1–242. doi: 10.3310/hta20420

Herlocher, M. L., Carr, J., Ives, J., Elias, S., Truscon, R., Roberts, N., & Monto, A. S. (2002). Influenza virus carrying an R292K mutation in the neuraminidase gene is not transmitted in ferrets. Antiviral research, 54(2), 99–111. doi: https://doi.org/10.1016/S0166-3542(01)00214-5

Hirotsu, N., Sakaguchi, H., Sato, C., Ishibashi, T., Baba, K., Omoto, S., … Watanabe, A. (2019). Baloxavir marboxil in Japanese pediatric patients with influenza: safety and clinical and virologic outcomes. Clin Infect Dis. doi: 10.1093/cid/ciz908

Hurt, A. C., Ernest, J., Deng, Y. M., Iannello, P., Besselaar, T. G., Birch, C., … Barr, I. G. (2009). Emergence and spread of oseltamivir-resistant A(H1N1) influenza viruses in Oceania, South East Asia and South Africa. Antiviral research, 83(1), 90–93. doi: 10.1016/j.antiviral.2009.03.003

Hurt, A. C., Hardie, K., Wilson, N. J., Deng, Y. M., Osbourn, M., Leang, S. K., … Barr, I. G. (2012). Characteristics of a widespread community cluster of H275Y oseltamivir-resistant A(H1N1)pdm09 influenza in Australia. J Infect Dis, 206(2), 148–157. doi: 10.1093/infdis/jis337

Hussain, M., Galvin, H. D., Haw, T. Y., Nutsford, A. N., & Husain, M. (2017). Drug resistance in influenza A virus: the epidemiology and management. Infection and drug resistance, 10, 121–134. doi: 10.2147/IDR.S105473

Imai, M., Yamashita, M., Sakai-Tagawa, Y., Iwatsuki-Horimoto, K., Kiso, M., Murakami, J., … Kawaoka, Y. (2019). Influenza A variants with reduced susceptibility to baloxavir isolated from Japanese patients are fit and transmit through respiratory droplets. Nature Microbiology. doi: 10.1038/s41564-019-0609-0

Ison, M. G., Portsmouth, S., Yoshida, Y., Shishido, T., Mitchener, M., Tsuchiya, K., … Hayden, F. G. (2020). Early treatment with baloxavir marboxil in high-risk adolescent and adult outpatients with uncomplicated influenza (CAPSTONE-2): a randomised, placebo-controlled, phase 3 trial. Lancet Infect Dis. doi: 10.1016/S1473-3099(20)30004-9

Ives, J. A., Carr, J. A., Mendel, D. B., Tai, C. Y., Lambkin, R., Kelly, L., … Roberts, N. A. (2002). The H274Y mutation in the influenza A/H1N1 neuraminidase active site following oseltamivir phosphate treatment leave virus severely compromised both in vitro and in vivo. Antiviral research, 55(2), 307–317. doi: 10.1016/s0166-3542(02)00053-0

Jones, J. C., Pascua, P. N. Q., Fabrizio, T. P., Marathe, B. M., Seiler, P., Barman, S., … Govorkova, E. A. (2020). Influenza A and B viruses with reduced baloxavir susceptibility display attenuated in vitro fitness but retain ferret transmissibility. Proceedings of the National Academy of Sciences, 201916825. doi: 10.1073/pnas.1916825117

Koszalka, P., Farrukee, R., Mifsud, E., Vijaykrishna, D., & Hurt, A. C. (2020). A rapid pyrosequencing assay for the molecular detection of influenza viruses with reduced baloxavir susceptibility due to PA/I38X amino acid substitutions. Influenza Other Respir Viruses. doi: 10.1111/irv.12725

Lackenby, A., Besselaar, T. G., Daniels, R. S., Fry, A., Gregory, V., Gubareva, L. V., … Meijer, A. (2018). Global update on the susceptibility of human influenza viruses to neuraminidase inhibitors and status of novel antivirals, 2016-2017. Antiviral research, 157, 38–46. doi: 10.1016/j.antiviral.2018.07.001

Li, T. C., Chan, M. C., & Lee, N. (2015). Clinical Implications of Antiviral Resistance in Influenza. Viruses, 7(9), 4929–4944. doi: 10.3390/v7092850

Meijer, A., Lackenby, A., Hungnes, O., Lina, B., van-der-Werf, S., Schweiger, B., … Zambon, M. (2009). Oseltamivir-resistant influenza virus A (H1N1), Europe, 2007-08 season. Emerg Infect Dis, 15(4), 552–560. doi: 10.3201/eid1504.181280

Mishin, V. P., Patel, M. C., Chesnokov, A., De La Cruz, J., Nguyen, H. T., Lollis, L., … Gubareva, L. V. (2019). Susceptibility of Influenza A, B, C, and D Viruses to Baloxavir. Emerg Infect Dis, 25(10), 1969–1972. doi: 10.3201/eid2510.190607

Noshi, T., Kitano, M., Taniguchi, K., Yamamoto, A., Omoto, S., Baba, K., … Naito, A. (2018). In vitro characterization of baloxavir acid, a first-in-class cap-dependent endonuclease inhibitor of the influenza virus polymerase PA subunit. Antiviral research. doi: 10.1016/j.antiviral.2018.10.008

Oh, D. Y., Barr, I. G., & Hurt, A. C. (2015). A Novel Video Tracking Method to Evaluate the Effect of Influenza Infection and Antiviral Treatment on Ferret Activity. PLOS ONE, 10(3), e0118780. doi: 10.1371/journal.pone.0118780

Omoto, S., Speranzini, V., Hashimoto, T., Noshi, T., Yamaguchi, H., Kawai, M., … Cusack, S. (2018). Characterization of influenza virus variants induced by treatment with the endonuclease inhibitor baloxavir marboxil. Sci Rep, 8(1), 9633. doi: 10.1038/s41598-018-27890-4

Patel, T. S., Cinti, S., Sun, D., Li, S., Luo, R., Wen, B., … Stevenson, J. G. (2017). Oseltamivir for pandemic influenza preparation: Maximizing the use of an existing stockpile. Am J Infect Control, 45(3), 303–305. doi: 10.1016/j.ajic.2016.09.024

Simonsen, L., Viboud, C., Grenfell, B. T., Dushoff, J., Jennings, L., Smit, M., … Holmes, E. C. (2007). The genesis and spread of reassortment human influenza A/H3N2 viruses conferring adamantane resistance. Mol Biol Evol, 24(8), 1811–1820. doi: 10.1093/molbev/msm103

Takashita, E., Daniels, R. S., Fujisaki, S., Gregory, V., Gubareva, L. V., Huang, W., … Meijer, A. (2020). Global update on the susceptibilities of human influenza viruses to neuraminidase inhibitors and the cap-dependent endonuclease inhibitor baloxavir, 2017-2018. Antiviral research, 175, 104718. doi: 10.1016/j.antiviral.2020.104718

Takashita, E., Ejima, M., Itoh, R., Miura, M., Ohnishi, A., Nishimura, H., … Tashiro, M. (2014). A community cluster of influenza A(H1N1)pdm09 virus exhibiting cross-resistance to oseltamivir and peramivir in Japan, November to December 2013. Euro Surveill, 19(1). doi: 10.2807/1560-7917.es2014.19.1.20666

Takashita, E., Ichikawa, M., Morita, H., Ogawa, R., Fujisaki, S., Shirakura, M., … Odagiri, T. (2019). Human-to-Human Transmission of Influenza A(H3N2) Virus with Reduced Susceptibility to Baloxavir, Japan, February 2019. Emerg Infect Dis, 25(11), 2108–2111. doi: 10.3201/eid2511.190757

Takashita, E., Kawakami, C., Ogawa, R., Morita, H., Fujisaki, S., Shirakura, M., … Odagiri, T. (2019). Influenza A(H3N2) virus exhibiting reduced susceptibility to baloxavir due to a polymerase acidic subunit I38T substitution detected from a hospitalised child without prior baloxavir treatment, Japan, January 2019. Euro Surveill, 24(12), 1900170. doi: 10.2807/1560-7917.ES.2019.24.12.1900170

Takashita, E., Kiso, M., Fujisaki, S., Yokoyama, M., Nakamura, K., Shirakura, M., … Tashiro, M. (2015). Characterization of a large cluster of influenza A(H1N1)pdm09 viruses cross-resistant to oseltamivir and peramivir during the 2013-2014 influenza season in Japan. Antimicrob Agents Chemother, 59(5), 2607–2617. doi: 10.1128/aac.04836-14

Tu, V., Abed, Y., Barbeau, X., Carbonneau, J., Fage, C., Lagüe, P., & Boivin, G. (2017). The I427T neuraminidase (NA) substitution, located outside the NA active site of an influenza A(H1N1)pdm09 variant with reduced susceptibility to NA inhibitors, alters NA properties and impairs viral fitness. Antiviral research, 137, 6–13. doi: https://doi.org/10.1016/j.antiviral.2016.11.007

Uehara, T., Hayden, F. G., Kawaguchi, K., Omoto, S., Hurt, A. C., De Jong, M. D., … Kida, H. (2019). Treatment-Emergent Influenza Variant Viruses With Reduced Baloxavir Susceptibility: Impact on Clinical and Virologic Outcomes in Uncomplicated Influenza. The Journal of Infectious Diseases. doi: 10.1093/infdis/jiz244

WHO. (2020). Recommended composition of influenza virus vaccines for use in the 2020– 2021 Northern Hemisphere influenza season. Retrieved from https://www.who.int/influenza/vaccines/virus/recommendations/202002_recommendation.pdf

Yen, H.-L., Herlocher, L. M., Hoffmann, E., Matrosovich, M. N., Monto, A. S., Webster, R. G., & Govorkova, E. A. (2005). Neuraminidase Inhibitor-Resistant Influenza Viruses May Differ Substantially in Fitness and Transmissibility. Antimicrobial Agents and Chemotherapy, 49(10), 4075–4084. doi: 10.1128/aac.49.10.4075-4084.2005

Zhou, B., Donnelly, M. E., Scholes, D. T., St George, K., Hatta, M., Kawaoka, Y., & Wentworth, D. E. (2009). Single-reaction genomic amplification accelerates sequencing and vaccine production for classical and Swine origin human influenza a viruses. J Virol, 83(19), 10309–10313. doi: 10.1128/JVI.01109-09

